# A DNA deliverer-receiver mechanism for DNA recruitment in phase-separated transcriptional condensates

**DOI:** 10.64898/2026.03.09.710715

**Authors:** Samuel Blazquez, Masataka Yamauchi, Tsuyoshi Terakawa

## Abstract

DNA transcription is a complex process involving numerous components that can assemble into phase-separated transcriptional condensates. However, whether condensates formed by multiple transcription factors behave through simple additive effects or instead exhibit non-additive, emergent properties remains unclear. Here, we use large-scale molecular dynamics simulations to investigate how three core transcription factors regulating pluripotency and early embryonic development—Nanog, Oct4, and Sox2—organize biomolecular condensates in the absence and presence of DNA. We find that condensate formation is primarily driven by intrinsically-disordered-region-mediated interactions of Nanog and Sox2, each of which individually promotes Oct4 phase separation; by contrast, when Nanog and Sox2 coexist in the absence of DNA, Oct4 is less efficiently incorporated into condensates. In the presence of DNA, condensates display a distinct spatial organization: Nanog and Sox2 form dense, well-mixed clusters, whereas Oct4 remains more dispersed in interstitial regions where DNA preferentially localizes, resulting in ∼20% higher DNA content in Oct4-containing condensates. Notably, phase separation reshapes DNA-protein interaction landscapes, altering the intramolecular regions that engage DNA. Together, these results support a synergistic DNA deliverer-receiver mechanism and suggest that non-additive, multi-component condensate organization constitutes an additional layer of gene expression regulation beyond canonical transcription factor-DNA binding.

## INTRODUCTION

Transcription factors (TFs) play a central role in gene regulation by recognizing specific DNA sequences and orchestrating the expression of target genes. Recent work shows that TFs and co-activators can assemble into condensates that promote gene activation. At super-enhancers, TFs assemble into condensates, which are enriched by RNA polymerase II (Pol II), chromatin architectural proteins like BRD4, and the Mediator complexes, through phase separation mediated by their intrinsically disordered regions (IDRs), giving rise to transcriptional condensates^1,2^. Pol II recruitment to transcriptional condensates has also been reconstituted *in vitro*^1,3,4^, demonstrating that these assemblies can concentrate the transcriptional machinery. In this context, condensate nucleation at enhancers can amplify gene expression through positive feedback among interacting components, supporting the view that condensates are functional microreactors for transcription rather than passive aggregates^5,6^.

Transcriptional control in pluripotent stem cells is not executed by a lone factor, but rather by an interconnected network of master regulators^7^. For example, Sox2, Oct4, and Nanog co-occupy many genomic loci, regulate one another’s expression, and together maintain the self-renewing, undifferentiated state of embryonic stem cells^8–11^. Each of these TFs contains intrinsically disordered regions (IDRs) within their sequences, which have been reported to endow them with the capacity to undergo phase separation and form condensates^12,13^. Single-molecule imaging and biophysical assays have shown that Sox2 forms DNA–protein co-condensates whose assembly depends on its intrinsically disordered region^2,14^. Although Oct4 alone does not form condensates, it undergoes phase separation when combined with the intrinsically disordered region of MED1, a core subunit of the Mediator complex^2,15^, and recent studies have proposed that Oct4 forms small clusters that promote DNA accessiblity^16^. Nanog forms gel-like condensates through its prion-like C-terminal domain that self-associate at low nanomolar concentrations and bridge DNA, and both coarse-grained molecular dynamics simulations and *in vitro* experiments reveal micelle-like internal organization within these Nanog condensates^17,18^. Despite extensive mapping of co-occupancy among Oct4, Sox2, and Nanog across the genome^19^, the molecular principles by which multiple TFs jointly assemble into condensates and how their interactions modulate DNA-binding specificity remain largely unknown. In particular, accumulating evidence suggests that the ability of TFs to form condensates cannot be explained by a simple additive contribution of individual components^20,21^. Also, both the molecular mechanisms, their interplay with DNA binding, and the biological significance of such non-additive contributions of each component in condensate formation remain largely unknown.

Experimental approaches have faced limitations in disentangling the individual interactions among TFs, or in resolving their co-localization with DNA, making it challenging to delineate the molecular basis of multicomponent condensate formation. To overcome this limitation, molecular dynamics simulations^22^, especially with amino-acid-resolution coarse-grained models^21,23–30^, have been employed, enabling quantitative and sequence-dependent analyses of multivalent interactions that drive phase separation. To date, coarse-grained molecular dynamics (CGMD) simulations of condensates containing Sox2, Oct4, and Nanog have been performed using the amino-acid-resolution AICG2+ model^31^ to represent intramolecular interactions and the HPS model^25^, combined with the Debye-Hückel potential for electrostatics, to describe intermolecular interactions responsible for phase separation^32^. The AICG2+ model has been combined with the 3SPN framework^33^, including its derivative 3SPN.2C model^34^, which represent DNA at a three-particle-per-nucleotide resolution, and together they have been successfully applied to elucidate the molecular mechanisms of various biomolecular systems^35–41^. With the implementation of GPU acceleration in biomolecular simulator OpenCafeMol, it has become possible to perform large-scale simulations^42^. In this study, we employed CGMD simulations to reconstruct the condensate formation of Sox2, Oct4, Nanog, and DNA, and quantitatively analyzed both the combination-dependent principles of condensate assembly and the molecular interactions between the condensates and DNA.

Our simulations revealed that Oct4 alone is inefficient at forming condensates but can do so in the presence of either Sox2 or Nanog. When Sox2 and Nanog coexist, however, Oct4 is excluded from the condensates, because both Oct4 and Nanog compete for binding to the same region in IDR of Sox2, indicating that condensate formation is not simply additive among components. We further found that Oct4 recruits DNA into condensates, whereas Nanog interferes with this recruitment, suggesting a functional asymmetry among TFs within multicomponent assemblies. In this context, Oct4 acts as a *deliverer* that captures and transports DNA, while Nanog- or Sox2-driven condensates function as receivers that provide a stable scaffold for DNA accumulation. Sox2 binds DNA nonspecifically through both its IDR and DNA-binding domain, but upon condensate formation, the IDR becomes engaged in IDR–IDR interactions, thereby biasing DNA association toward the DNA-binding domain. Together, these findings demonstrate that multicomponent condensates go beyond concentration-dependent accumulation of TFs, providing a potential regulatory mode in which specific factor combinations in a condensate modulate DNA co-localization and binding specificity.

## RESULTS

### Deciphering the intermolecular interactions within TF condensates

To elucidate the interactions operating within TF condensates, we selected three TFs that play essential roles in development and cellular homeostasis: Nanog, Oct4, and Sox2. Nanog is crucial for maintaining pluripotency and self-renewal in embryonic stem cells^43^. Oct4 is a key DNA-binding protein that functions as a pioneer factor, facilitating chromatin opening and DNA recruitment^44,45^. Sox2, often acting in synergy with Oct4^46,47^, is essential for lineage specification and early embryonic development^48–50^. Although a variety of experimental techniques—such as fluorescence recovery after photobleaching (FRAP), GFP fluorescence recovery, coalescence assays, and microrheology—have been employed to address the intermolecular forces underlying biomolecular condensates^51–54^, molecular dynamics (MD) simulations provide a powerful complementary tool by enabling residue-and domain- level interrogation of interaction networks and their dynamic reorganization within condensates. They have been successfully used to predict the multiphase architectures formed by protein-protein aggregation^55^, as well as the complex internal structures and interfacial properties of biomolecular condensates^56^ or their viscoelastic properties^57,58^. Experiments and simulations have also been synergistically combined to clarify salt-dependent phase behavior, in which condensate formation is driven by electrostatic interactions at low salt, suppressed at intermediate salt concentrations, and re-emerges at high salt due to hydrophobic and other non-ionic interactions^59^. In this work, we explore molecular interactions using MD simulations, starting from pre-assembled condensate states (Fig. 1a) of binary or ternary mixtures of the three TFs mentioned above. This strategy allows us to efficiently access condensate regimes, while avoiding the prohibitively long simulation times that would be required to observe spontaneous phase separation.

**Figure 1:**
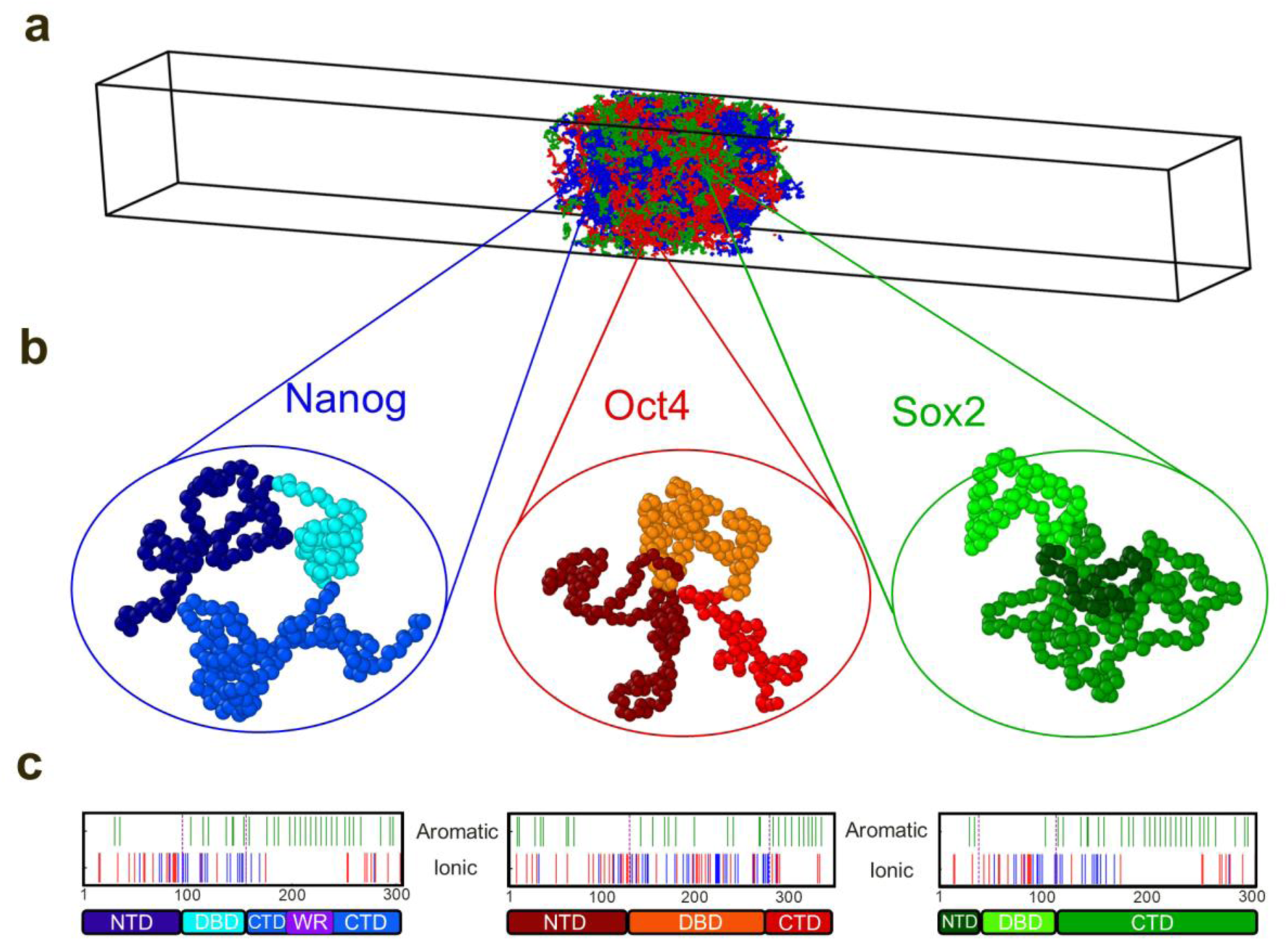
Coarse-grained molecular dynamics simulations of phase-separated TFs. **(a)** Representative initial snapshot of the simulation system showing a phase-separated condensate composed of Nanog, Oct4, and Sox2 within a confined simulation box. **(b)** Coarse-grained representations of the three TFs analyzed in this study: Nanog (left), Oct4 (center), and Sox2 (right). Beads are colored according to structural domains. For Nanog: N-terminal domain (NTD, dark blue), DNA-binding domain (DBD, cyan), and C-terminal domain (CTD, light blue). For Oct4: NTD (dark red), DBD (orange), and CTD (light red). For Sox2: NTD (forest green), DBD (light green), and CTD (dark green). **(c)** Sequence features of Nanog, Oct4, and Sox2. The upper panels indicate residue-level chemical properties: aromatic residues (F, Y, W) in green, negatively charged residues (D, E) in red, and positively charged residues (K, R) in blue. The lower panels show domain organization, color-coded as in (b), including NTD, DBD, and CTD. In Nanog, a tryptophan-rich repeat region (WR) is additionally highlighted (purple).

The TFs studied in this work were represented using a simplified molecular model in which each amino-acid residue is described by a single bead located at its C_α_ atom position (Fig. 1b). All three TFs contain intrinsically disordered N-terminal (NTD) and C-terminal (CTD) domains [in the case of Nanog, the CTD also includes a tryptophan-repeated (WR) region], which were modeled using the flexible local potentials, which defines bond-angle and dihedral-angle terms based on statistical distributions of loop regions in folded proteins^60^. In contrast, the structured DNA-binding domain (DBD) was modeled with the AICG2+^31,60^ potential, which stabilizes the reference structure while capturing native fluctuations. Intermolecular interactions include electrostatic, hydrophobic, and excluded-volume interactions, where electrostatics are described by the Debye-Hückel model and hydrophobic interactions are captured by the HPS model^61^.

Generally, the main regions that govern biomolecular condensation are intrinsically disordered domains enriched in aromatic and charged residues, which give rise to π–π and cation–π interactions characteristic of proteins prone to condensate formation^62,63^. Taking this into account, we examined the chemical composition of the TFs studied in this work (Fig. 1c), highlighting aromatic residues (green), cationic residues (blue), and anionic residues (red) along with their structural domains. This analysis shows that, in all three proteins, aromatic residues are predominantly located in CTD, while positively charged amino acids are concentrated in DBD. In the cases of Nanog and Sox2, the DBD is relatively short, and the CTD accounts for a large portion of the protein length. By contrast, Oct4 has a markedly longer DBD and a relatively short CTD. These differences in CTD length—being the domain richest in aromatic residues—are expected to be critical for understanding the distinct phase separation behaviors of these TFs, as will be discussed later.

To decipher the intermolecular interactions responsible for condensate formation among Nanog, Oct4, and Sox2, we began by simulating a mixture containing 67 molecules of each TF, forming a phase-separated condensate placed in the center of an elongated simulation box (Fig. 2a, top, Supplementary Movie 1). We performed simulations of 10⁸ MD steps, corresponding to approximately 1 μs in physical time. To assess equilibration, we monitored the time evolution of the *z*-positions of all proteins (Fig. 2b). Starting from artificially compressed initial configurations (See Methods), Nanog and Sox2 relaxed into stable, condensate-like assemblies that remained equilibrated throughout the second half of the simulation. In contrast, Oct4 displayed a markedly lower tendency to form condensate—as previously reported^32^—yet the presence of Nanog and Sox2 modestly increased the fraction of Oct4 residing within the condensate (Fig. 2a, bottom, Supplementary Movie 1). This trend was not observed in previous simulations of Oct4 alone, using the same models and shorter simulations, showing how this TF by itself was not able to form condensate^32^.

**Figure 2:**
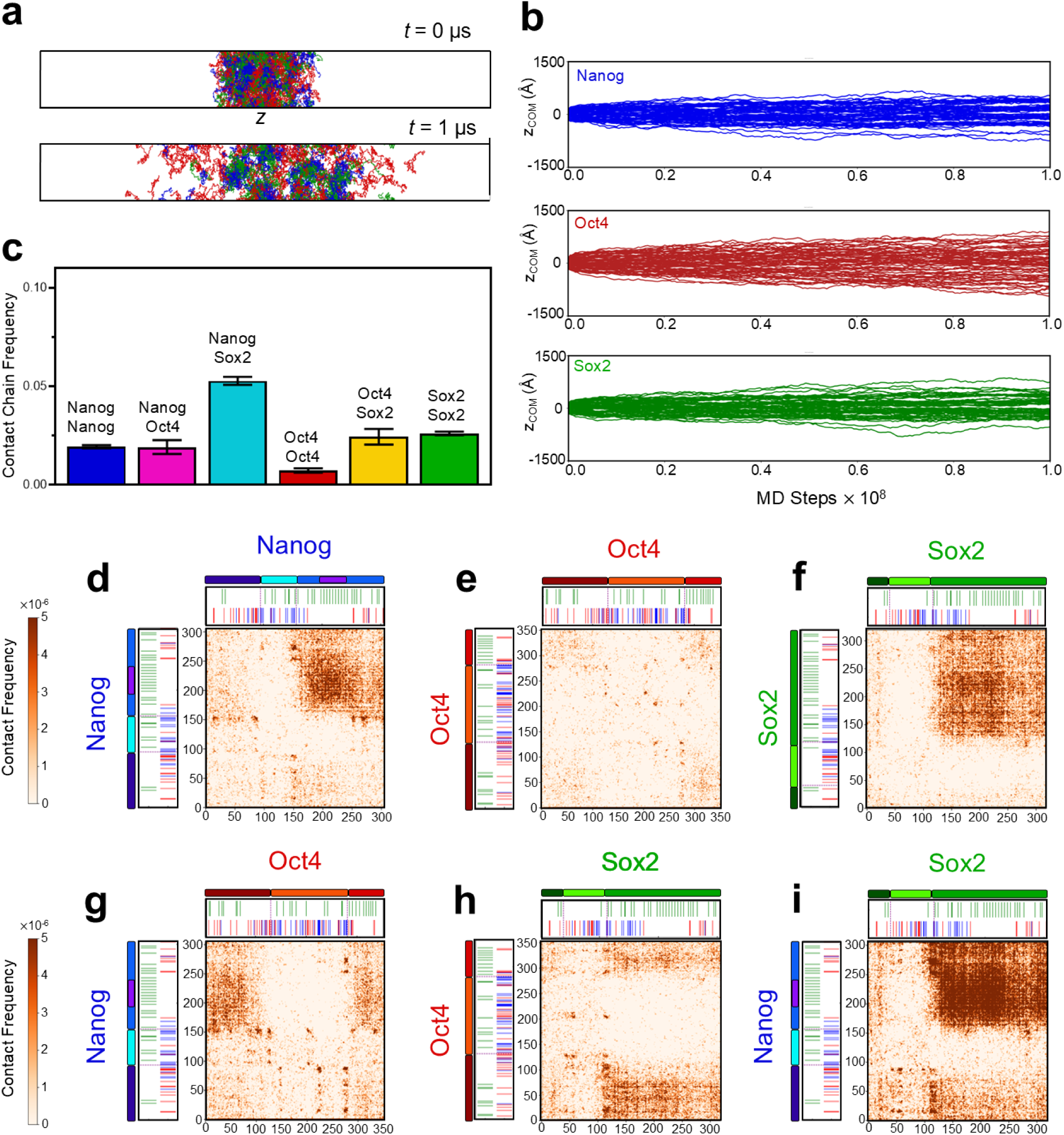
Nanog, Oct4, and Sox2 form phase-separated condensates driven by CTD-mediated interactions. **(a)** Representative snapshots of the three-component simulation slab at *t* = 0 μs (top) and *t* = 1 μs (bottom), illustrating condensate formation along the *z*-axis. **(b)** Time evolution of the center-of-mass positions (*z*_COM_) for 67 Nanog (blue, top), 67 Oct4 (red, middle), and 67 Sox2 (green, bottom) molecules during the simulation, showing spatial confinement within the condensed phase. **(c)** Intermolecular contact frequencies in the Nanog-Oct4-Sox2 mixture. Bars indicate pairwise chain contacts, color-coded according to the interacting species. **(d–i)** Residue-level contact maps for homotypic and heterotypic interactions: (d) Nanog-Nanog, (e) Oct4-Oct4, (f) Sox2-Sox2, (g) Nanog-Oct4, (h) Oct4-Sox2, and (i) Nanog-Sox2. Contact probabilities are shown as heat maps. For each panel, the domain architecture and sequence chemical features of the corresponding proteins are displayed above and to the left of the map. Domain and chemical feature coloring follows Fig. 1.

To elucidate the molecular interactions driving condensate formation among the TFs, we first examined their homotypic and heterotypic interactions—recently shown to play a key role in condensate formation^64^. To this end, we quantified the chain–chain contact frequencies (Fig. 2c). These data reveal that Nanog–Sox2 interactions overwhelmingly dominate the multicomponent condensate, relegating all other TF–TF interactions to secondary roles and almost completely suppressing Oct4–Oct4 contacts. The final snapshot supports this behavior, showing that Nanog sequesters Sox2 while largely excluding Oct4 from the condensate (Fig. 2a, bottom). Among the remaining interactions, Sox2–Sox2 and Oct4–Sox2 contacts are the next most frequent, although far less prominent than Nanog–Sox2. Altogether, the three-component mixture is clearly governed by heterotypic interactions.

We next mapped the residue–residue interactions for all TF pairs, as described in Methods. Nanog–Nanog contacts are concentrated in the CTD (Fig. 2d), highlighting the importance of this intrinsically disordered, aromatic-rich region. Importantly, the Nanog CTD also mediates interactions with Oct4: both the NTD and CTD of Oct4 contact the Nanog CTD (Fig. 2g). In contrast, Oct4–Oct4 interactions are minimal and consist mainly of isolated electrostatic contacts between dispersed charged residues (Fig. 2e).

For Sox2, CTD once again emerges as the principal interaction hub. The strongest Sox2–Sox2 contacts (Fig. 2f) and the Oct4–Sox2 interactions (Fig. 2h) are both mediated by the Sox2 CTD. Finally, the Nanog–Sox2 contact map (Fig. 2i) confirms that their interactions are driven by the CTDs of both proteins, representing the strongest heterotypic contacts among all pairs analyzed. Taken together, these results demonstrate that in the Nanog–Oct4–Sox2 mixture, condensate formation is governed primarily by heterotypic CTD–CTD interactions between Nanog and Sox2.

We next sought to examined whether the TF–TF interactions change upon removal of the third TF, testing whether its presence in the ternary mixture alters the binary interactions between the remaining TFs. To address this, we performed simulations starting from pre-compressed configurations, in which a binary mixture of Nanog–Oct4, Oct4–Sox2, or Nanog–Sox2 was placed at the center of an elongated simulation box. Each system contained 100 molecules of each TF (200 proteins in total). As before, we monitored the positions of the proteins along the z-axis over time. For Nanog–Oct4, the trajectories show that Nanog forms a much more stable and denser condensate, reaching equilibrium around 6×10⁷ steps and remaining stable thereafter (Fig. 3a). Oct4, in contrast, displays a clear tendency to remain more dispersed along the z-axis—although its distribution also stabilizes after 6×10⁷ steps—consistent with its limited intrinsic condensate-forming propensity. This behavior is evident in the final snapshot of the simulation (Fig. 3d, Supplementary Movie 2). Chain–chain contact frequencies confirm this trend: Nanog–Nanog interactions are markedly stronger than both Oct4–Oct4 and, to a lesser degree, Nanog–Oct4 contacts (Fig. 3j). Consequently, the resulting condensate is predominantly Nanog-rich, with heterotypic interactions mainly serving to recruit Oct4 into the cluster. As observed previously, the interior architecture adopts a micelle-like organization^32^ (Fig. 3g), with the CTD (blue) buried inside and the NTD/DBD (dark blue and cyan) facing the exterior.

**Figure 3:**
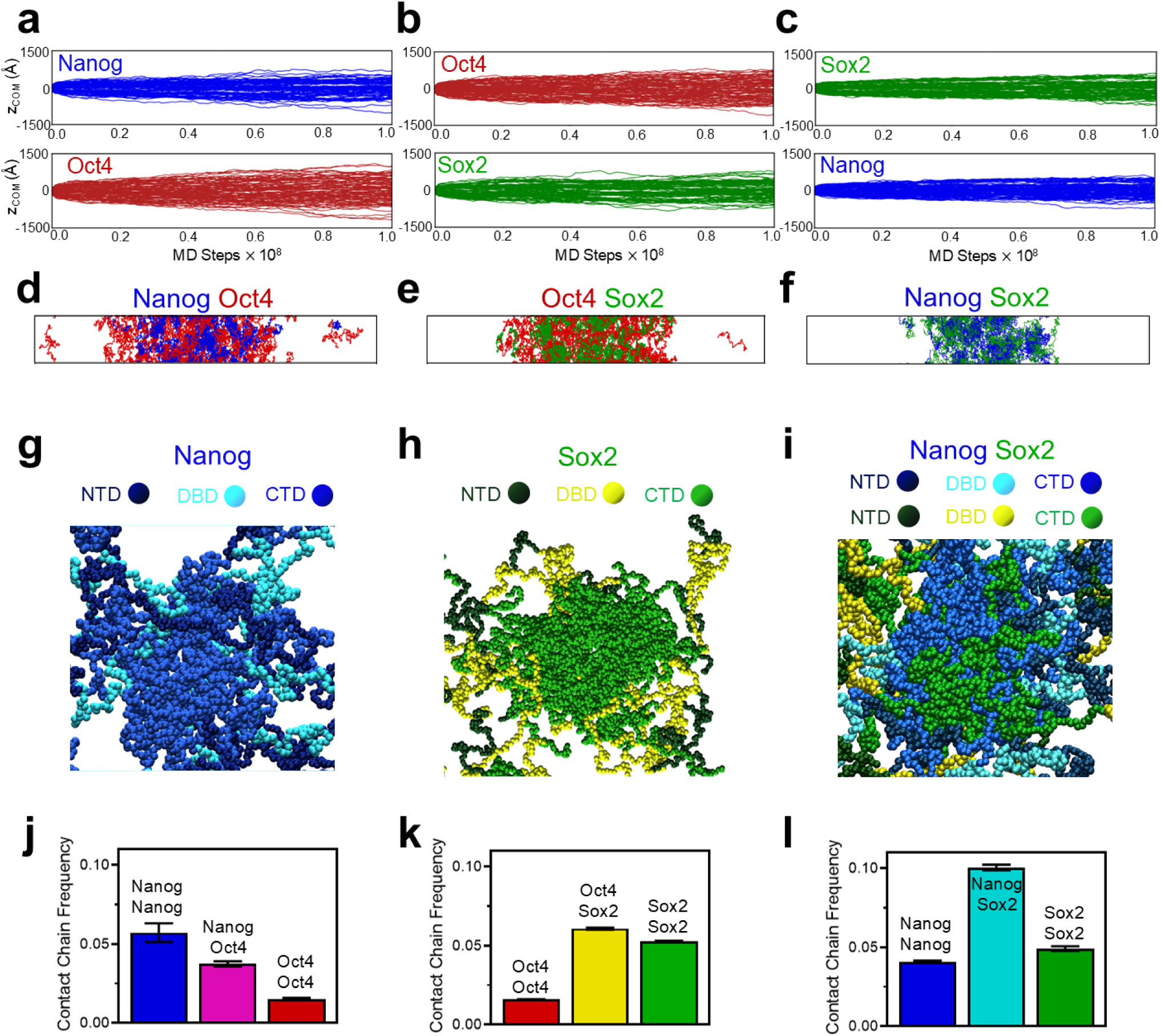
Binary mixture simulations highlight the central roles of Nanog and Sox2 in promoting phase separation. (a–c) Time evolution of the center-of-mass positions (*z*_COM_) of TFs along the slab normal (*z*-axis) in binary mixtures: (a) 100 Nanog (blue, top) and 100 Oct4 (red, bottom); (b) 100 Oct4 (red, top) and 100 Sox2 (green, bottom); and (c) 100 Sox2 (green, top) and 100 Nanog (blue, bottom). **(d–f)** Final snapshots of the binary simulations showing condensate formation in (d) Nanog–Oct4, (e) Oct4–Sox2, and (f) Nanog–Sox2 mixtures. **(g–i)** Representative close-up views of condensates. (g) Nanog–Nanog condensate colored by structural domains: NTD (dark blue), DBD (cyan), and CTD (light blue). (h) Sox2–Sox2 condensate colored by structural domains: NTD (dark green), DBD (yellow), and CTD (light green). (i) Nanog–Sox2 condensate colored following the same coding as (g-h). Across systems, condensates exhibit a micelle-like organization, with CTDs preferentially localized in the interior and NTDs/DBDs enriched at the periphery. **(j–l)** Intermolecular contact frequencies in the binary mixtures: (j) Nanog–Oct4, (k) Oct4–Sox2, and (l) Nanog–Sox2. Bars indicate pairwise chain contacts, color-coded according to the interacting species.

Distinct behavior emerged when Oct4 was mixed with Sox2. Surprisingly, Oct4 was less dispersed (Fig. 3b), than in the Nanog–Oct4 system, and a larger fraction of Oct4 molecules remained embedded in the condensate (Fig. 3e, Supplementary Movie 3). Contact frequency analysis revealed that Oct4–Sox2 interactions were the strongest in this system (Fig. 3k), supporting experimental evidence of cooperativity between these TFs^46,47,65,66^. Oct4 self-interactions, in contrast, play only a minor role. As with Nanog, Sox2 robustly recruited Oct4 into the condensate—an effect even stronger than in the Nanog–Oct4 mixture. Interestingly, this trend contrasts with the ternary mixture, where Nanog–Sox2 interactions dominate, and Oct4–Sox2 cooperativity is weakened probably due to Nanog sequestering Sox2. Importantly, the condensate architecture of Sox2 shows a similar micelle structure as found for Nanog, with the inner CTD (light green) and outer DBD (yellow) and NTD (dark green) as shown in Fig. 3h.

Finally, simulations of the Nanog–Sox2 system revealed, consistent with the ternary mixture, the strongest contacts among all binary combinations (Fig. 3l). The protein distributions along the *z*-axis showed both TFs tightly localized at the center of the simulation slab, forming the densest and most stable condensate (Fig. 3c), as also observed in the final simulation snapshot (Fig. 3f, Supplementary Movie 4). Owing to the favorable heterotypic interactions, the resulting condensates were well mixed, exhibiting the same micelle-like architecture described above, with CTDs forming the condensate core and NTD/DBD regions facing the exterior (Fig. 3i). Nonetheless, homogeneous Nanog–Nanog or Sox2–Sox2 clusters also appeared.

Taken together, these binary simulations (performed in the absence of DNA) reinforced the central role of the CTD—and, to a lesser extent, the NTD—in driving condensate formation, while the DBD appeared to play only a minor role, if any. Importantly, the residue–residue contact maps (Supplementary Fig. 1) displayed no qualitative differences from those in the ternary system: Nanog–Nanog and Sox2–Sox2 interactions were governed by their CTDs, and Oct4 interacted through both NTD and CTD regions. Crucially, both Nanog and Sox2 effectively recruited Oct4 into condensates, whereas Oct4 alone showed a very limited propensity for condensate formation. Finally, the comparison between the binary and ternary systems highlighted the delicate interplay among the three TFs: Nanog disrupted Oct4–Sox2 cooperativity by sequestering Sox2, thereby largely excluding Oct4 from the condensate.

### Three-to-One TF-DNA simulations

Following the characterization of the molecular interactions driving condensate formation for Nanog, Oct4, and Sox2, we next investigated DNA–protein interactions. Before simulating a complete phase-separated transcriptional condensate, we examined DNA binding through three-to-one TF-DNA simulations to isolate the intrinsic DNA affinity of each TF, which would otherwise be masked by condensate formation. These simulations consisted of one DNA molecule in the presence of single molecules of Nanog, Oct4, and Sox2 proteins. Interactions between DNA and proteins were primarily nonspecific and included excluded-volume and electrostatic contributions.

We first computed the distance between DNA and each protein as a function of time for three independent trajectories (Fig. 4a). These results showed that, on average, the three TFs remained within 8–15 Å of the DNA throughout the simulations. Plotting the averaged DNA–protein distances for the three TFs (Fig. 4b) revealed that binding is not simultaneous: at certain moments one or two TFs detach from the DNA while the remaining one(s) stay bound. This indicated that DNA–protein binding is highly dynamic, with periods in which individual TFs move closer to or further from the DNA. Representative snapshots were shown in Fig. 4c: (i) a configuration where all three TFs simultaneously contact the DNA, and examples where only one TF is bound — Nanog (ii), Oct4 (iii), or Sox2 (iv).

**Figure 4:**
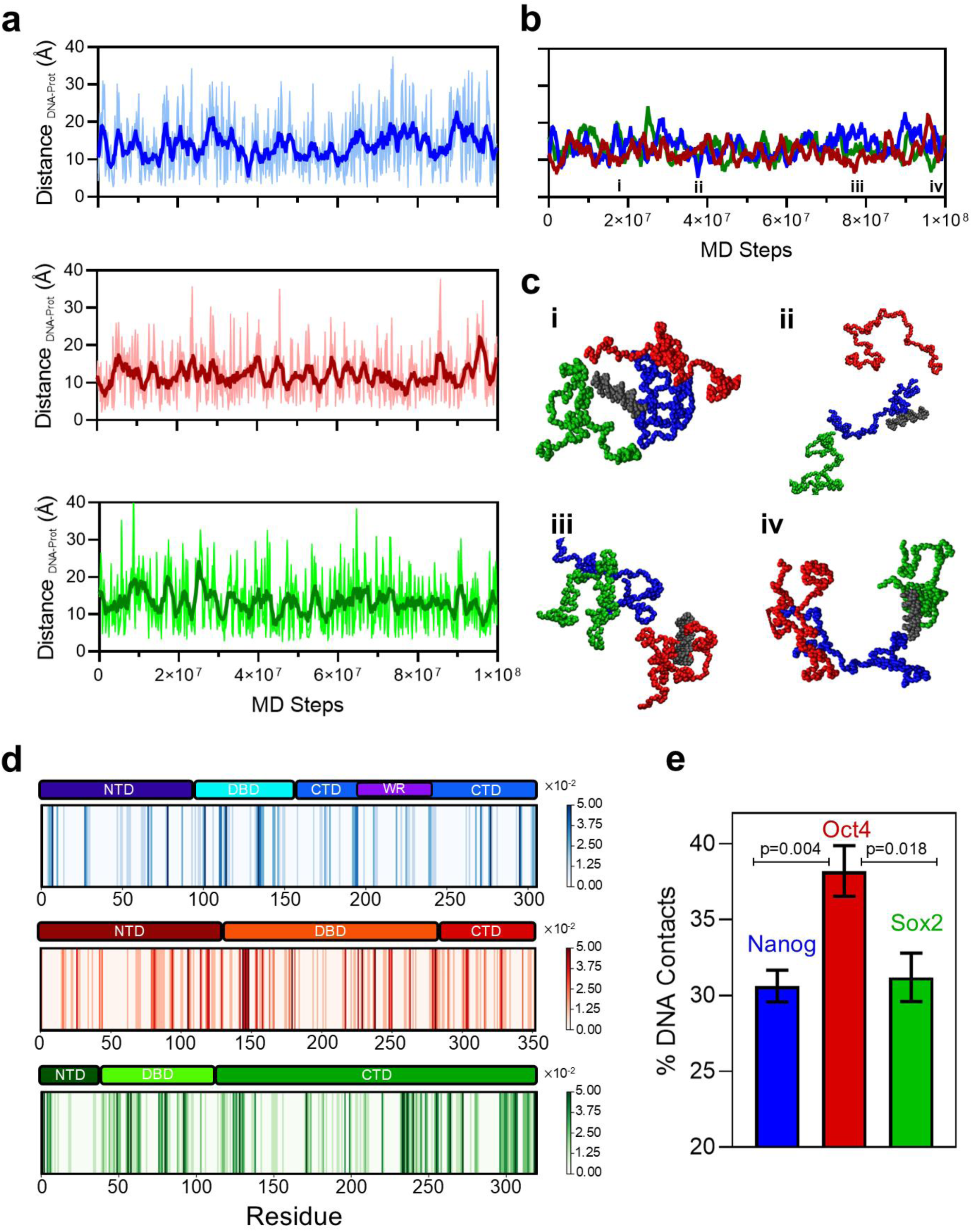
Single-molecule simulations reveal enhanced DNA affinity of Oct4. **(a)** Time evolution of the minimum distance between DNA and each TF. Bold traces indicate the mean values averaged over three independent simulations, and shaded regions represent the corresponding deviations. Top: Nanog (blue). Middle: Oct4 (red). Bottom: Sox2 (green). **(b)** Direct comparison of the averaged DNA–protein distances for Nanog (blue), Oct4 (red), and Sox2 (green) plotted together. **(c)** Representative snapshots from single-molecule simulations showing DNA (gray) interacting with Nanog (blue), Oct4 (red), and Sox2 (green). Panels i–iv correspond to the time points indicated in (b). **(d)** Residue-level DNA contact maps for Nanog (top), Oct4 (middle), and Sox2 (bottom). Structural domains are indicated above each map. **(e)** Quantification of relative DNA affinity derived from simulations containing DNA and all three TFs. Bars represent the percentage of DNA contacts formed by Nanog (blue), Oct4 (red), and Sox2 (green). Statistical significance was assessed using paired t-tests with Holm–Šídák correction for multiple comparisons (α = 0.05).

Next, we mapped contacts between each amino acid bead of the proteins and any of the DNA beads (Fig. 4d). For Nanog (Fig. 4d, top), the strongest and most numerous interactions were found in the DBD and surrounding regions, mainly due to electrostatic interactions between the negatively charged DNA backbone phosphates and the abundance of cationic residues. Interactions involving the NTD and CTD were also observed, though less prominently. In the case of Oct4 (Fig. 4d, middle), DNA interactions were distributed across the protein sequence, although the strongest interactions were again located in the DBD, which is enriched in cationic residues. Finally, the DNA–Sox2 contact profile (Fig. 4d, bottom) revealed that, although the DBD displayed high affinity for DNA, the CTD also emerged as a region of unexpectedly strong DNA binding—at least in the absence of condensate formation, as will be discussed later.

Nevertheless, mapping intermolecular interactions alone did not allow us to quantify DNA–protein affinity. To address this issue, we quantified, for each TF, the fraction of simulation frames in which the distance between the center of mass of DNA and the protein were less than 10 Å over 10^8^ MD steps simulations for three independent seeds. Our simulations revealed that Oct4 exhibits a markedly and statistically significantly higher affinity for DNA (about 39 ± 2 %) compared to Nanog (31 ± 1 %) and Sox2 (about 31 ± 2 %), which display almost identical DNA affinities (Fig. 4e). Taken together, these findings demonstrated that, at the single-molecule level, Oct4 has an enhanced DNA affinity relative to Nanog and Sox2.

### Architecture of Phase-Separated Transcriptional Condensates

Up to this point, we focused on simulations of three-to-one systems, in which three different TF—Nanog, Oct4, and Sox2—were mixed with a single DNA molecule. To investigate how condensate formation influences TF–DNA binding, we next performed simulations of TF–DNA condensates containing multiple copies of both TFs and DNA. In these simulations, condensates were composed of 67 molecules of each TF in the ternary mixtures (or 100 molecules of each TF for the binary mixtures) and 80 double-stranded DNA molecules, each 20 bp in length, with a poly(dC–dG) sequence. Systems were simulated either as binary mixtures (Nanog–Oct4, Oct4–Sox2, or Nanog–Sox2) or as ternary mixtures containing all three TFs. The use of short DNA fragments was necessitated by technical constraints associated with periodic boundary conditions, which preclude the inclusion of much longer DNA molecules without introducing spurious self-interactions. Nevertheless, these short DNA segments are sufficient to capture the local features of TF–DNA interactions, thereby providing a reasonable model for assessing how condensate formation modulates DNA association. The simulations were initiated from mixed TF–DNA configurations that were artificially compressed and subsequently allowed to relax, enabling the formation and reorganization of condensate-like TF–DNA assemblies.

We first analyzed the condensate structures from representative simulation snapshots of the Nanog–Oct4 (Fig. 5a), Oct4–Sox2 (Fig. 5b), and Nanog–Sox2 (Fig. 5c) binary mixtures, as well as the Nanog–Oct4–Sox2 ternary mixture (Fig. 5d, Supplementary Movie 5). In the first two binary mixtures (Nanog–Oct4 and Oct4–Sox2), the TFs with condensate-forming propensity—Nanog and Sox2—formed strongly self-associated, stable condensates, while Oct4 localized around these clusters with markedly fewer inter- and intra-protein associations. These results showed that the condensates were not uniformly mixed. Although some interactions occurred between different TFs, especially in the Oct4–Sox2 mixture, Nanog and Sox2 mainly clustered with their own kind instead of with Oct4. The higher DNA affinity of Oct4 revealed by the three-to-one TF–DNA simulations was also clearly reflected in the snapshots: DNA molecules avoided the dense Nanog–Nanog or Sox2–Sox2 clusters—which remained largely inaccessible—and instead localized in the surrounding regions enriched in Oct4. The third binary mixture, Nanog–Sox2, displayed different behavior. Unlike Oct4-containing systems, these condensates were well mixed, and DNA appeared to associate equally with both TFs. This agreed with the three-to-one TF–DNA simulations (Fig. 4e), which revealed very similar DNA affinities for Nanog and Sox2. Finally, the ternary mixture in the presence of DNA again showed a well-mixed Nanog–Sox2 condensate depleted in DNA, while DNA predominantly localized near Oct4, as observed in the binary mixtures containing Oct4.

**Figure 5:**
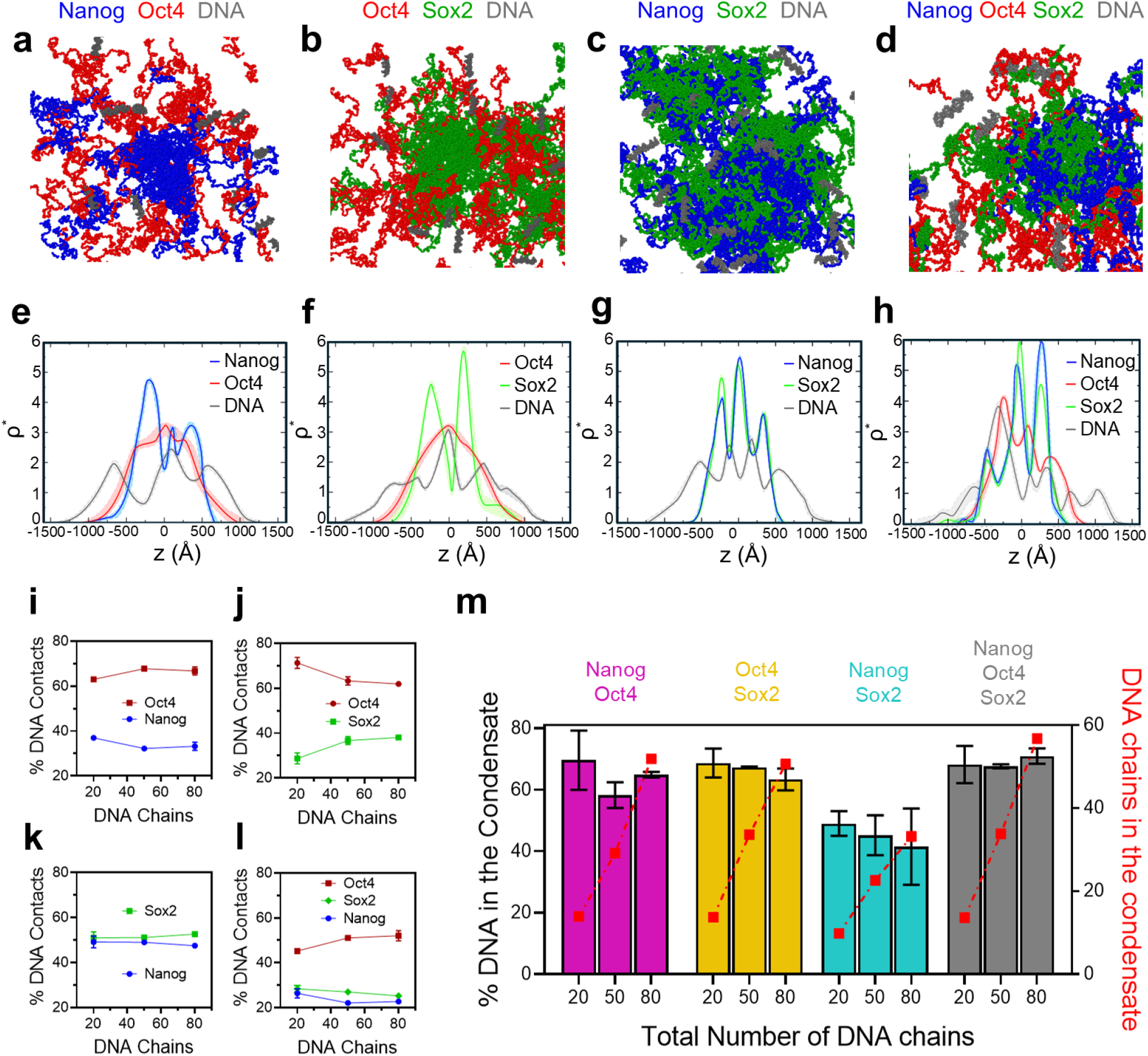
Structural organization of Nanog, Oct4, and Sox2 condensates at varying DNA concentrations. (a-d) Representative snapshots of phase-separated condensates formed in the presence of 80 DNA molecules: (a) Nanog-Oct4, (b) Oct4-Sox2, (c) Nanog-Sox2, and (d) Nanog-Oct4-Sox2 mixtures. **(e-h)** One-dimensional density profiles along the simulation box axis for Nanog (blue), Oct4 (red), Sox2 (green), and DNA (gray). Systems include (e) 100 Nanog + 100 Oct4 + 80 DNA, (f) 100 Oct4 + 100 Sox2 + 80 DNA, (g) 100 Nanog + 100 Sox2 + 80 DNA, and (h) 67 Nanog + 67 Oct4 + 67 Sox2 + 80 DNA. Shaded regions represent the uncertainty of the measurements. **(i–l)** Relative DNA-binding affinities of TFs as a function of increasing DNA concentration in (i) Nanog-Oct4, (j) Oct4-Sox2, (k) Nanog-Sox2, and (l) Nanog-Oct4-Sox2 mixtures. Error bars, when not visible, are smaller than the symbol size. **(m)** Quantification of DNA partitioning into condensates. Colored bars (left y-axis) show the percentage of total DNA localized within the condensate, while red squares (right y-axis) indicate the corresponding number of DNA chains inside the condensed phase. Results are shown for Nanog-Oct4 (magenta), Oct4-Sox2 (mustard), Nanog-Sox2 (cyan), and Nanog-Oct4-Sox2 (gray) mixtures as a function of the total number of DNA chains in the simulation.

To quantify these architectural features, we computed density profiles along the z-axis of the simulation box. We used the reduced density (*ρ\**) as a general measure, defined as the density of beads of a component divided by its expected density under a uniform distribution. Values of *ρ\** > 1 indicated enrichment and condensate formation, whereas values < 1 indicated depletion. This *ρ\** threshold (*ρ\** > 1) was also used to define the condensate boundaries for the calculation of critical concentrations and DNA content. The density profiles of the Nanog–Oct4 (Fig. 5e) and Oct4–Sox2 (Fig. 5f) mixtures in the presence of 80 DNA molecules were consistent with the snapshots: regions enriched in Nanog or Sox2 were depleted in DNA, whereas DNA localized in Oct4-rich regions. Oct4 showed broader and less intense density peaks, reflecting its lower tendency to form compact clusters. Similar behavior was observed at other DNA concentrations (Supplementary Fig. 6a and b). In contrast, the Nanog–Sox2 mixture with 80 DNA molecules (Fig. 5g) showed perfectly overlapping and narrow density peaks, confirming a well-mixed condensate. Interestingly, in this case, the DNA distribution depended on concentration: at low DNA levels, DNA was randomly dispersed within the condensate (Supplementary Fig. 6c), whereas at high concentrations, DNA became depleted from the condensate interior and accumulated in regions not occupied by Nanog–Sox2 clusters. This reflected the strong Nanog–Sox2 interactions combined with the relatively weaker DNA–TF affinities. Finally, the ternary mixture (Fig. 5h) exhibited features consistent with the binary simulations: overlapping Nanog and Sox2 profiles indicating a well-mixed scaffold, enrichment of Oct4 in DNA-accessible regions, and preferential localization of DNA in Oct4-rich zones.

We next quantified DNA affinity as a function of DNA concentration. In Oct4-containing binary mixtures (Fig. 5i–j), Oct4 consistently showed higher DNA affinity than Nanog or Sox2. By contrast, Nanog–Sox2 mixtures showed nearly identical DNA affinities for both TFs (∼50%; Fig. 5k), although with a slightly larger affinity for Sox2. The ternary mixture (Fig. 5l) followed the same trend: Oct4 exhibited the highest DNA affinity, while Nanog and Sox2 bound DNA similarly. DNA affinity showed only minimal and non-monotonic dependence on DNA concentration in all cases. Importantly, this affinity increased upon condensate formation: while three-to-one TF–DNA simulations yielded Oct4–DNA affinity of 39%, this value increased to approximately 50% within condensates. Moreover, comparison between DNA affinities obtained in two-to-one TF-DNA simulations (see Methods) led to similar conclusions showing affinities of 57% (Nanog–Oct4) and 60% (Oct4–Sox2) which increased to 65% in binary condensate simulations. Thus, condensate formation amplifies the relative DNA affinity of Oct4. While DNA affinity captures the strength of TF–DNA association, it does not describe how DNA is spatially distributed within the condensate. We therefore quantified the fraction of DNA located inside the condensate for all four mixtures (Fig. 5m). Condensates containing Oct4 consistently incorporated ∼70% of the DNA across concentrations, whereas the Nanog–Sox2 condensate incorporated only ∼50%. In the Nanog–Sox2 system, the DNA fraction decreased as DNA concentration increased. We further quantify the absolute number of DNA chains contained within it (Fig. 5m, red dashed line). This analysis indicates that, within the tested range, the number of DNA chains inside the condensate does not reach saturation. Increasing the initial number of DNA chains leads to a proportional increase in the absolute number of chains within the condensate, even when the corresponding fractions remain comparable.

### DNA-Protein interactions change upon condensate formation

With the architecture and composition of the transcriptional condensates established up to this point, we next focused on the molecular protein–protein and protein–DNA interactions within condensates formed in the presence of DNA. We primarily analyzed the ternary mixture of the three TFs with 80 DNA molecules, although results for binary mixtures and other DNA concentrations were provided in the Supplementary Material and referenced throughout this section.

Mapping protein–protein interactions revealed that the presence of DNA did not substantially alter the main intermolecular contacts within the condensates. As observed previously, the CTDs of Nanog and Sox2 remained the dominant contributors to Nanog–Nanog (Fig. 6a) and Sox2–Sox2 (Fig. 6c) interactions, whereas Oct4–Oct4 interactions remained weak (Fig. 6b), even in the presence of DNA. In contrast, Oct4 interacted with both Nanog (Fig. 6d) and Sox2 (Fig. 6e) through its NTD and CTD. Consistent with earlier observations, Oct4–Sox2 interactions were stronger than Nanog–Oct4 interactions, reinforcing the synergistic relationship between these proteins, particularly in the presence of DNA. Moreover, Nanog–Sox2 interactions (Fig. 6f), mediated by the CTDs of both proteins, were the strongest heterotypic interactions within the condensate. These interaction patterns agreed with the structural features observed above (Fig. 5a–d), where Nanog–Oct4 and Oct4–Sox2 formed non-homogeneous assemblies, while Nanog–Sox2 formed well-mixed condensates characterized by strong heterotypic contacts. Additionally, the same conclusions were found when analyzing the protein–protein contact maps of the binary mixtures at different DNA concentrations (Supplementary Figs. 2–4).

**Figure 6:**
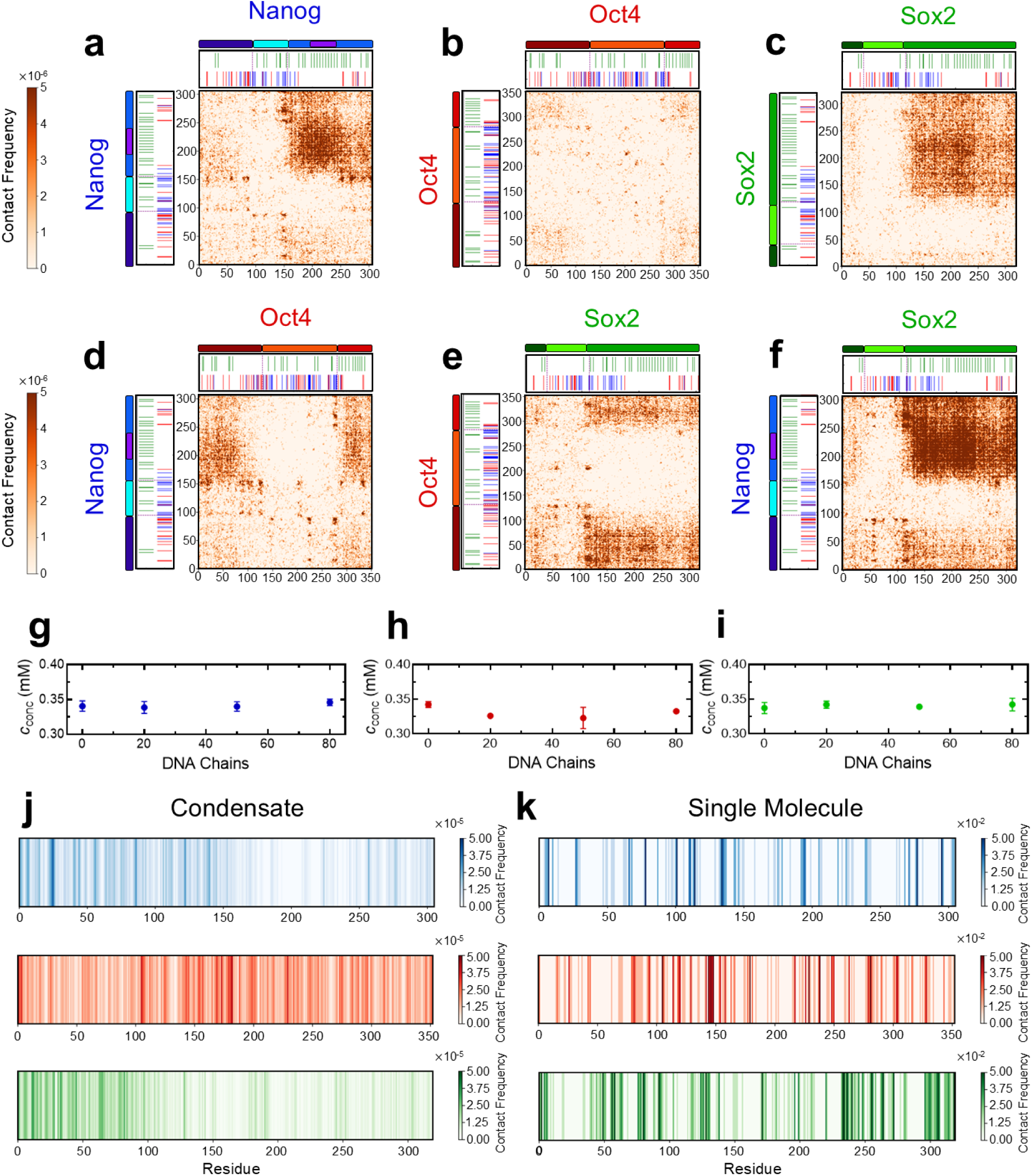
Molecular interaction patterns of Nanog, Oct4, and Sox2 within DNA-containing phase-separated condensates. (a-f) Residue-level protein–protein contact maps obtained from simulations containing 67 Nanog, 67 Oct4, 67 Sox2, and 80 DNA chains. Structural domains and sequence chemical features are displayed above and alongside each map with the same color coding as in Fig. 1. (a) Nanog-Nanog; (b) Oct4-Oct4; (c) Sox2-Sox2; (d) Nanog-Oct4; (e) Oct4-Sox2; (f) Nanog-Sox2 interactions. **(g-i)** Critical concentrations in the condensed phase as a function of DNA chain number for (g) Nanog, (h) Oct4, and (i) Sox2. **(j)** Residue-resolved DNA–protein contact frequencies within the phase-separated condensate (67 Nanog, 67 Oct4, 67 Sox2, 80 DNA). Top to bottom: Nanog-DNA, Oct4-DNA, and Sox2-DNA. **(k)** DNA-protein contact frequencies from single-molecule simulations containing one copy each of Nanog, Oct4, Sox2, and DNA. Top to bottom: Nanog-DNA, Oct4-DNA, and Sox2-DNA.

We also examined the critical concentrations of the condensed phases for each component, calculated using density profiles (Fig. 5 and Supplementary Fig. 6) and defined by ρ* > 1 as previously mentioned. Across all tested DNA concentrations (20, 50, and 80 DNA molecules), DNA showed no measurable effect on the critical concentrations of the condensed phases, which remained approximately constant at ∼0.34 mM within error bars for the three studied TFs: Nanog, Oct4, and Sox2 (Fig. 6g–i). Thus, within the conditions explored here, DNA neither enhanced nor suppressed condensate formation, and protein–protein interactions remained essentially unchanged. These conclusions are also observed for binary mixtures of all TFs as shown in Supplementary Figs. 7–9. We note that variations in DNA length or the inclusion of specific protein–DNA interactions could alter this behavior, although these effects are beyond the scope of the present work.

In contrast to protein–protein interactions and critical concentrations, condensate formation had a pronounced impact on protein–DNA interactions. Upon condensation, the CTDs of Nanog and Sox2 became engaged in stabilizing the condensate, rendering them largely inaccessible to DNA. Consequently, DNA interactions with the Nanog CTD observed in single-molecule simulations of the three TFs together in presence of one molecule of DNA (Fig. 6k, top), weakened or disappeared in the condensate, with binding shifting primarily to the NTD and DBD at high DNA concentration (Fig. 6j, top). This effect was not only found in the ternary mixture, but similar behavior was also observed in the Nanog–Oct4 binary mixture condensate simulations at all studied concentrations (Supplementary Fig. 5a) and in the comparison with the contact maps of single molecule simulations of the binary mixture (one Nanog, one Oct4, and one DNA) showed in Supplementary Fig 7b. The effect was even more pronounced for Sox2: while three-to-one TF–DNA simulations showed strong DNA interactions with the Sox2 CTD (Fig. 6k, bottom), these contacts almost vanished in condensates, with DNA binding redirected mainly to the NTD at high DNA concentration (Fig. 6j, bottom), a trend also observed in the Oct4–Sox2 and Nanog–Sox2 binary mixtures condensate simulations across all DNA concentrations studied (Supplementary Fig. 5b–c) and which is compared with the single molecule simulations of binary mixtures in Supplementary Figs. 8b and 9b. In sharp contrast, Oct4–DNA interactions remained largely unchanged between single-molecule (Fig. 6k, middle) and condensate simulations (Fig. 6j, middle), involving multiple binding regions—most prominently the DBD. This behavior reflected Oct4’s lower intrinsic condensate-forming propensity: as discussed above, Oct4 was retained within the condensate primarily through interactions with Nanog or Sox2 rather than through strong self-association, leaving it comparatively free to engage with DNA.

## DISCUSSION

In recent years, condensate formation primarily via liquid–liquid phase separation had become a subject of enormous interest. In this work, our simulations supported the recently proposed hypothesis of phase-separated transcriptional condensates, in which TFs, together with coactivators, formed clusters and recruited promoters and enhancers within these assemblies^2,67^. We focused on three fundamental TFs intimately linked to pluripotency and gene regulation—Nanog, Oct4, and Sox2. Our simulations showed the strong tendency of Nanog and Sox2 to undergo phase separation, as well as their role in enhancing and enabling the phase separation of Oct4 when in contact with them. Moreover, we provided further support for experimental evidence demonstrating the cooperative behavior between Oct4 and Sox^46,47,66,68^. Finally, we showed that when the three TFs were mixed, Nanog sequestered Sox2, nearly suppressing Oct4-Sox2 interactions and thereby adding further complexity to the previously well-established cooperativity between Oct4 and Sox2. Importantly, this behavior could not be inferred from pairwise interactions alone, indicating that condensate formation gives rise to intrinsically non-additive effects among TFs.

We elucidated the molecular interactions driving condensate formation in Nanog, Oct4, and Sox2 mixtures, showing that both in the absence and presence of DNA at different concentrations, CTD-mediated interactions dominated condensate formation. This was consistent with the literature, where IDRs were identified as the main regions responsible for condensate formation^69,70^, and the CTD represented the largest and most highly disordered domain in these TFs. These predominant CTD-driven interactions gave rise to micelle-like condensates, in which the CTDs were buried inside the condensate, while the DBDs and NTDs remained exposed on the outside—consistent with previous findings for Nanog–Nanog^18^ and, as revealed here, also for Nanog–Sox2 and Sox2–Sox2 clusters. Such micelle-like reorganization implies that the effective interaction landscape within condensates is not a linear superposition of individual TF–TF interactions, but instead emerges from collective, non-additive coupling mediated by condensate architecture.

Recent studies hypothesized a high affinity of Oct4 for DNA^44,71,72^. Here, we confirmed this DNA-delivering role through three-to-one TF–DNA simulations quantifying DNA–Oct4, DNA–Nanog, and DNA–Sox2 affinities in mixtures containing all three TFs. Our results revealed a significantly higher affinity of DNA for Oct4 compared to Nanog or Sox2, underscoring the importance of Oct4 in transcriptional condensates despite its limited intrinsic tendency to phase separate. These affinities were also reflected in the structural features of the relevant phase-separated transcriptional condensates. In binary mixtures of Nanog or Sox2 with Oct4, Nanog–Nanog and Sox2–Sox2 condensates were TF-rich and DNA-poor, while DNA preferentially associated with Oct4, which remained more dispersed within the condensate due to its lower condensate-forming propensity. In contrast, Nanog–Sox2 mixtures demonstrated that the DNA content within these condensates was markedly lower than when Oct4 was present, providing strong evidence for a condensate-specific biological role of Oct4 as a DNA deliverer. Notably, this condensate-specific function of Oct4 was not predictable from its intrinsic DNA affinity or from binary TF–DNA interactions, but instead emerged only in the context of multicomponent condensate formation, underscoring a fundamentally non-additive effect captured by our simulations. This effect was further supported by the ternary mixture simulations, in which the DNA content was similar to that found in the binary mixtures containing Oct4, due to the presence of Oct4.

Moreover, our simulations showed that DNA–Oct4 interactions were broadly dispersed along the protein, whereas DNA contacts with Nanog and Sox2 were concentrated mainly in the DBD and CTD (less prominently for Nanog). Notably, upon condensate formation, DNA interactions with Nanog—and especially with Sox2—shifted toward the NTD, reflecting the sequestration of their CTDs in stabilizing the condensates. These changes in the location of protein–DNA contacts revealed the impact of condensate formation on DNA–TF molecular interactions, emphasizing the importance of understanding these processes at the molecular level.

Our large-scale simulations of phase-separated transcriptional condensates reveal a molecular mechanism driven by the synergistic interplay between Nanog or Sox2 and Oct4. Nanog and Sox2 act as scaffolds that robustly promote condensate formation, whereas Oct4—despite its limited intrinsic propensity to phase separate on its own^32^—is retained within these condensates through interactions with its partners. Once incorporated, Oct4 assumes a central functional role as a DNA deliverer: it is more broadly distributed throughout the condensate and preferentially engages DNA. These findings led us to propose a deliverer–receiver mechanism (Fig. 7). In this model, Oct4 functions as the deliverer by capturing DNA from outside the condensate (delivered DNA, gray), enabled by its relatively high DNA affinity and low self-clustering tendency, and subsequently distributing it within the condensate. In contrast, Nanog and Sox2 function as receivers, forming stable and less mobile condensates that incorporate and retain the recruited DNA (received DNA, yellow). This framework provides a molecular-level explanation for how TFs coordinate protein–DNA interactions within phase-separated condensates, offering new insights into gene regulation and broader implications for biophysical and biomedical research. Importantly, our simulations predict that condensate formation generates non-additive effects on both protein–protein and protein–DNA interactions, leading to emergent functional roles that cannot be inferred from pairwise interactions alone.

**Figure 7:**
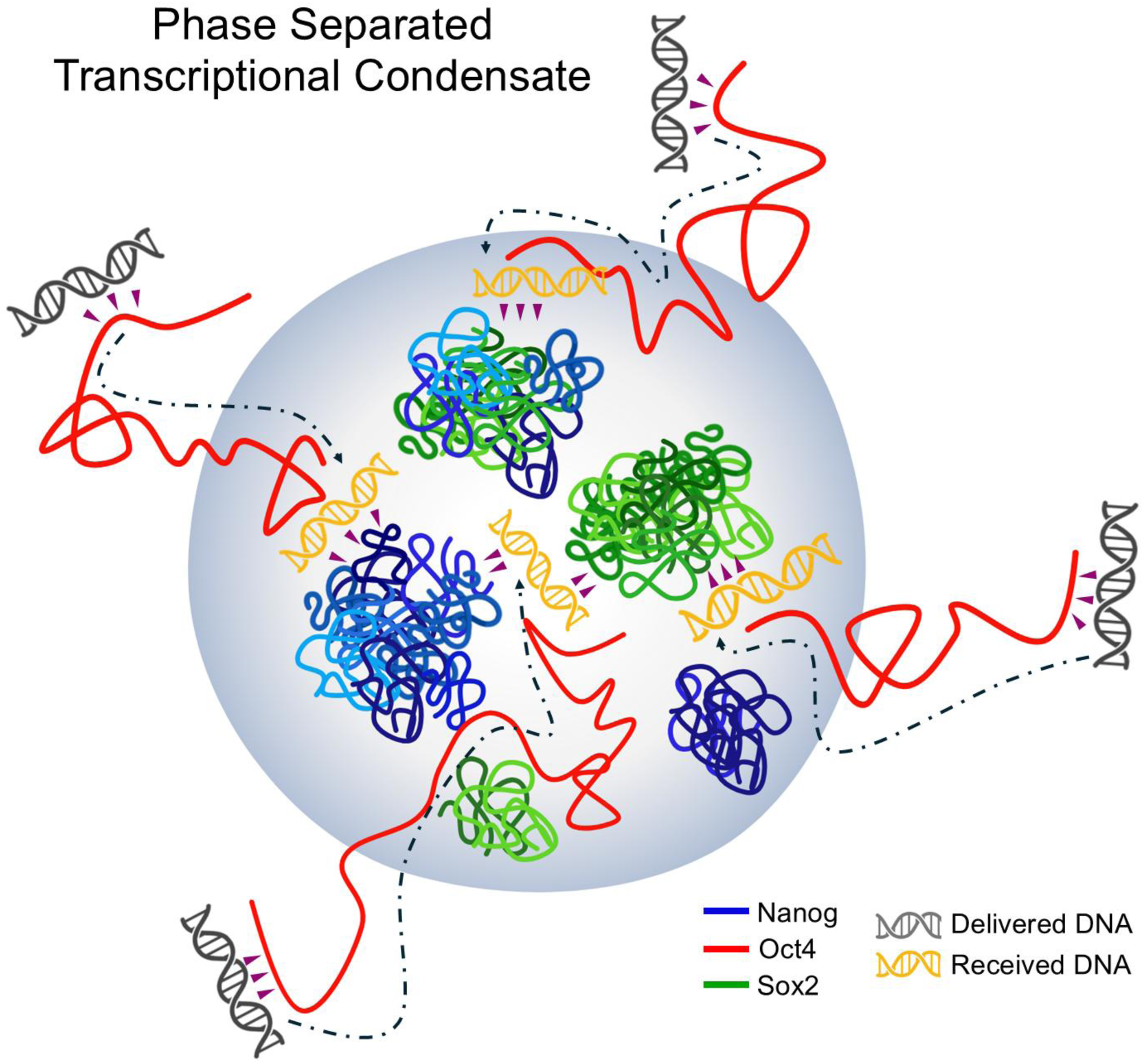
Proposed deliverer–receiver mechanism within phase-separated transcriptional condensates. Schematic model of the deliverer–receiver mechanism. Within the condensate (grey shaded region), Oct4 (red) is broadly dispersed and does not form stable self-clusters. In contrast, Nanog (blue) and Sox2 (green) assemble into relatively stable clusters, which may be homogeneous or mixed in composition. In this framework, Oct4 functions as a *deliverer* by capturing DNA molecules from outside the condensate (delivered DNA, grey) and facilitating their entry and redistribution within the condensed phase. The recruited DNA is subsequently incorporated into the more stable and less mobile Nanog/Sox2 clusters, which act as *receivers* (received DNA, yellow).

Such non-additive behavior provides a natural framework for understanding how transcriptional condensates can act as regulatory hubs that integrate multiple inputs into context-dependent transcriptional outcomes. In the context of pluripotency, small fluctuations in the relative abundance of Nanog, Sox2, and Oct4 could therefore result in switch-like changes in gene expression programs^73^, enabling robust maintenance of cell identity while preserving the capacity for rapid state transitions. Our simulations suggest that condensate formation amplifies these nonlinear effects by reorganizing protein–DNA interactions in a collective and emergent manner.

The predictions arising from our simulations can be directly tested by combining controlled perturbations of TF composition with quantitative readouts of DNA localization and transcriptional output. In particular, *in vitro* reconstituted condensate systems with tunable stoichiometries of Nanog, Sox2, and Oct4 would allow direct assessment of non-additive effects, for example by examining whether the addition or removal of Oct4 induces qualitative, rather than incremental, changes in DNA partitioning within condensates. Single-molecule fluorescence imaging or super-resolution microscopy could further resolve how DNA–TF contact patterns are reorganized upon condensate formation, while reporter-based transcription assays would enable correlation of these structural changes with functional transcriptional outcomes. In cellular contexts, systematic modulation of TF expression levels—using inducible or titratable systems—combined with live-cell imaging and nascent transcription measurements could test whether small changes in TF ratios produce switch-like or nonlinear responses in gene expression, as predicted by our simulations. Together, such approaches would provide a rigorous experimental framework to validate the emergent, non-additive regulatory roles of transcriptional condensates suggested by this study.

Despite the insights provided by our simulations, several limitations of this study should be acknowledged. First, our coarse-grained modeling approach necessarily simplifies molecular details, such as sequence-specific DNA recognition, post-translational modifications, and explicit solvent effects, which may influence TF behavior in vivo. Second, the DNA molecules used in the simulations were short fragments introduced for technical reasons related to periodic boundary conditions, and therefore do not capture long-range chromatin architecture, higher-order DNA topology, or nucleosome-associated constraints present in cellular genomes. Third, while our simulations systematically explored binary and ternary mixtures with defined stoichiometries, cellular transcriptional condensates are embedded in far more complex environments that include additional cofactors, chromatin-associated proteins, and active processes such as transcription and ATP-dependent remodeling. Finally, the timescales accessible to the simulations are limited, and thus our results primarily reflect steady-state or quasi-equilibrium behaviors rather than the full dynamics of condensate assembly and disassembly. In this context, although the simulations are consistent with the proposed ‘deliverer–receiver’ framework, the current data primarily capture steady-state organization and do not directly resolve the dynamic process by which DNA might be transferred between factors. For example, the behavior of Oct4 observed in the simulations may also be interpreted as acting in an anchor-like DNA-retaining role rather than directly demonstrating active DNA delivery. Future experimental approaches, such as single-molecule imaging, will therefore be important to directly test the proposed dynamic delivery process and to evaluate the mechanistic validity of the deliverer–receiver model. These limitations highlight the need for future studies that integrate more detailed molecular models, longer DNA substrates, and direct experimental validation to fully assess the biological relevance of the non-additive effects predicted here.

## MATERIALS AND METHODS

### Coarse-grained molecular models

We used the full-length sequences of three Mus musculus TFs: Nanog, Oct4, and Sox2. Each amino acid was represented as a single bead positioned at the Cα atom. All three TFs possessed intrinsically disordered regions at both their N- and C-termini (NTD and CTD). The specific domain boundaries for each protein are defined as follows: Nanog consists of an NTD (residues 1–95), a DBD (residues 96–155), and a CTD (residues 156–305). Oct4 comprises an NTD (residues 1–130), a DBD (residues 131–280), and a CTD (residues 281–352). Sox2 includes an NTD (residues 1–39), a DBD (residues 40–112), and a CTD (residues 113–319). The disordered domains were represented using flexible local potentials that imposed bond-angle and dihedral-angle restraints derived from statistical analyses of loop regions in folded proteins^60^. By contrast, the structured DNA-binding domain (DBD) was treated with the AICG2+^31,60^ potential, which preserved the native globular fold while allowed thermal fluctuations around the reference structure. Intermolecular interactions in the simulations accounted for electrostatic, hydrophobic, and excluded-volume effects: electrostatics were modeled using the Debye–Hückel formalism, whereas hydrophobic interactions were described by the HPS model^61^, a framework extensively employed in studies of intrinsically disordered proteins and liquid–liquid phase separation^74–77^. For DNA, each nucleotide was represented by three beads corresponding to the base, sugar, and phosphate groups. The 3SPN.2C potential^34^ was applied to preserve B-form geometry, reproduce sequence-dependent curvature, and accurately capture the persistence length and melting temperature of dsDNA. This protein–DNA combination of models had been rigorously validated in multiple studies^36–38,78,79^, establishing its robustness and reliability in faithfully capturing both structural fidelity and protein–DNA interaction behavior. DNA–protein interactions in the simulations were largely nonspecific in nature, arising from a combination of excluded-volume effects and electrostatic forces contributions. For intra- and inter-protein interactions, charged amino acids were assigned a unit charge. In the case of DNA, phosphate groups carried a charge of −0.6e for intra-DNA interactions, accounting for counterion condensation along the backbone. For protein–DNA interactions, however, phosphate charges were set to −1.0e to reflect counterion release upon complex formation.

Reference structures for the three TFs were obtained following recent work^32^. Briefly, we employed MODELLER^80^ to generate an all-atom structural model from the PDB entry (PDB ID: 2VI6^81^) for Nanog, which was used as the reference structure for this protein. In the case of Sox2 and Oct4, we used the structures predicted by AlphaFold2^82^. To generate the 20 bp dsDNA fragments, we employed the 3DNA software^83^.

### Molecular dynamics simulations

As in previous simulation studies of biomolecular condensates^18,32^, three independent condensate simulations were carried out for each system in a long rectangular box, with small X and Y dimensions and an extended Z dimension, under periodic boundary conditions applied in all three directions. The box side lengths were 320, 320, and 3000 Å along the X, Y, and Z axes, respectively. For the binary mixtures of TFs, 100 molecules of each TF were used, whereas for the ternary mixture, a total of 201 molecules were included (67 of each type). For simulations at different DNA concentrations, 20, 50, or 80 double-stranded DNA molecules (20 bp) with a poly(dC–dG) sequence, (CG)₁₀ each were included in the previously described systems. To prepare the initial phase-separated configurations, 67 molecules of each TF in the ternary mixture and 100 molecules of each TF in the binary mixtures (and the corresponding number of DNA molecules) were first placed on grids in compact conformations while avoiding overlaps. Short MD runs (of the order of 10^4^ and 10^5^ MD steps depending on the system) with anisotropic pressure of 5 bar for X and Y axis were then conducted while gradually decreasing the box size along the Z axis until a condensate of 320 × 320 × 320 Å³ size was formed. In all cases, after obtaining the condensate, this was equilibrated in the cubic box with a NVT simulation of 1 × 10^6^ MD steps.

For the three-to-one TF–DNA simulations, we employed a cubic box with side lengths of 120 Å in each dimension (120 × 120 × 120 Å³), and three TFs (one of each type) and one DNA molecule were included in the box. For the two-to-one TF-DNA simulations, we included one DNA molecule and two different TFs. The same coarse-grained models as in the condensate simulations were employed.

In all the simulations, Langevin dynamics was used to integrate the equations of motion with a timestep of 0.2 CafeMol time units (∼0.01 ps). Simulations were performed in the NVT ensemble at a monovalent salt concentration of 100 mM, which has been shown in previous studies to promote phase separation for these proteins^18,32^ and a temperature of 300 K, and were carried out using OpenCafeMol^42^. All simulations were run for 1 × 10⁸ MD steps, corresponding to approximately 1 µs.

In most analyses, we used the Python library MDAnalysis^84^ to read DCD files and analyze trajectories. To compute protein–protein contact maps in each condensate simulation, we counted the number of contacts between amino acids in every frame using a residue–residue cutoff distance of 6.5 Å, averaging over all frames and chains. For the chain–chain contact frequency, we calculated how many times one chain was in contact with another, either homotypic or heterotypic, using the same 6.5 Å cutoff. The number of observed contacts was then divided by the total number of possible contacts, given by *n_A_ × n_B_ × n_frames_*, where *n_A_* and *n_B_* denote the number of chains of types A and B, respectively, and *n_frames_* is the total number of frames in the simulation.

## ACKNOWLEDGEMENT

We would like to thank members of the Theoretical Biophysics laboratory at Kyoto University for discussions and assistance throughout this work. We are particularly grateful to Dr. Azuki Mizutani and Prof. Shoji Takada for generously providing the initial structures and configuration files for the molecular dynamics simulations. This work was supported by the Grant-in-Aid for Transformative Research Areas (24H00882 to T.T.), the grant from the Takeda Science Foundation (to T.T.), the grant from the Shimazu Science Foundation (to T.T.), the grant from the Yamada Science Foundation (to T.T.), and a Short-Term Postdoctoral Fellowship for Research in Japan of the JSPS (PE25017 to S.B.).

## DATA AVAILABILITY

The data that support the findings of this study are available within the article and Supplementary Information files.

---

**Supplementary Figure 1.**
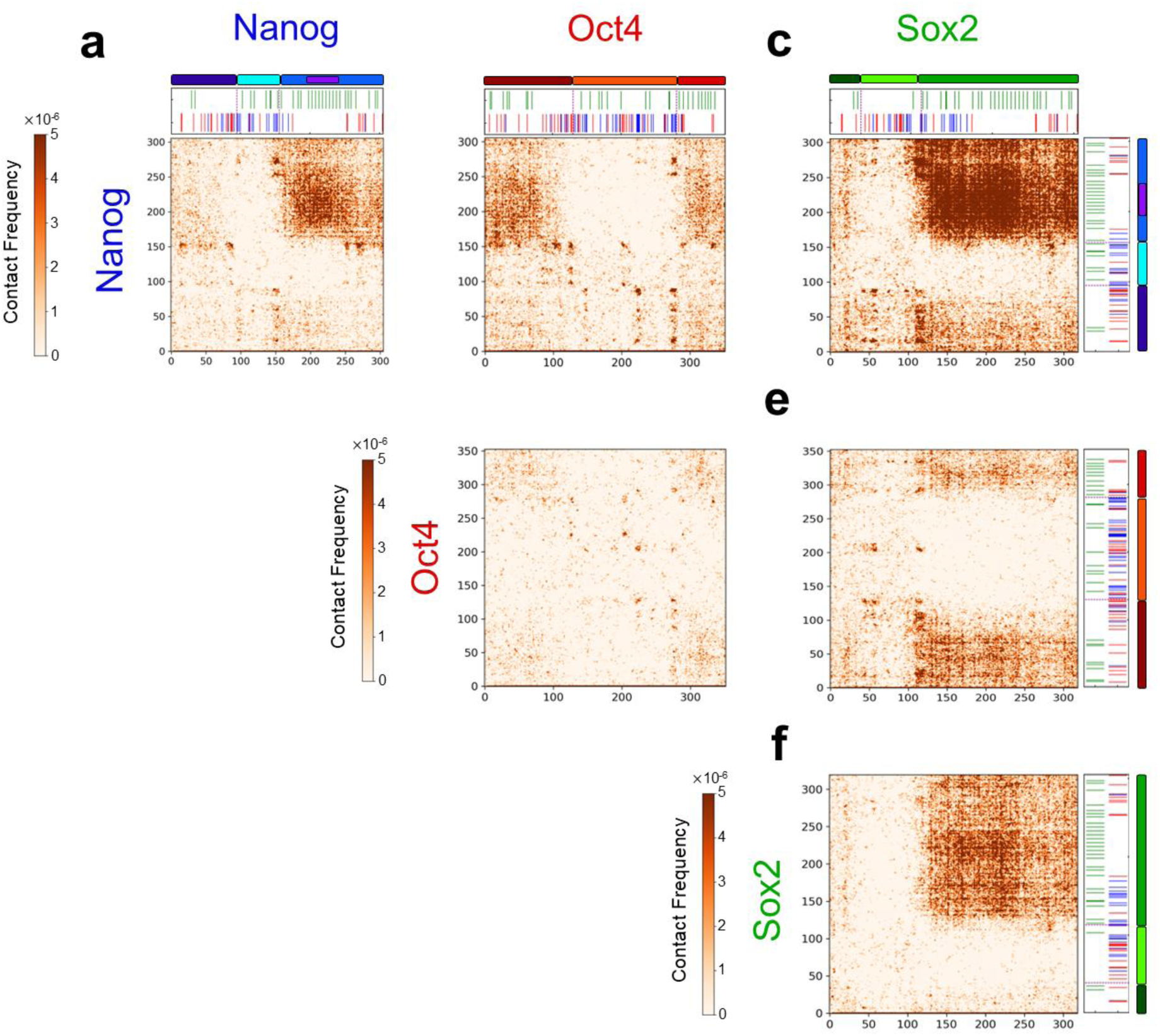
: Protein-protein interaction maps in binary TF condensates. **(a)** Nanog–Nanog residue-level contact map averaged over the Nanog-Oct4 and Nanog-Sox2 binary mixtures. The domain architecture and sequence chemical features of Nanog are shown above the map. **(b)** Nanog-Oct4 contact map derived from the Nanog-Oct4 binary system. Domain and sequence annotations are displayed as in (a). **(c)** Nanog-Sox2 contact map from the Nanog-Sox2 binary system. Structural domains and sequence features of Sox2 are shown above the map, and those of Nanog are indicated on the right. **(d)** Oct4-Oct4 contact map averaged over the Nanog-Oct4 and Oct4-Sox2 binary mixtures. **(e)** Oct4–Sox2 contact map from the Oct4-Sox2 binary system. Domain organization and sequence features of Oct4 are shown on the right. **(f)** Sox2-Sox2 contact map averaged over the Oct4-Sox2 and Nanog-Sox2 binary mixtures. Domain organization and sequence features of Sox2 are shown on the right following the same color code as in Fig. 1.

**Supplementary Figure 2.**
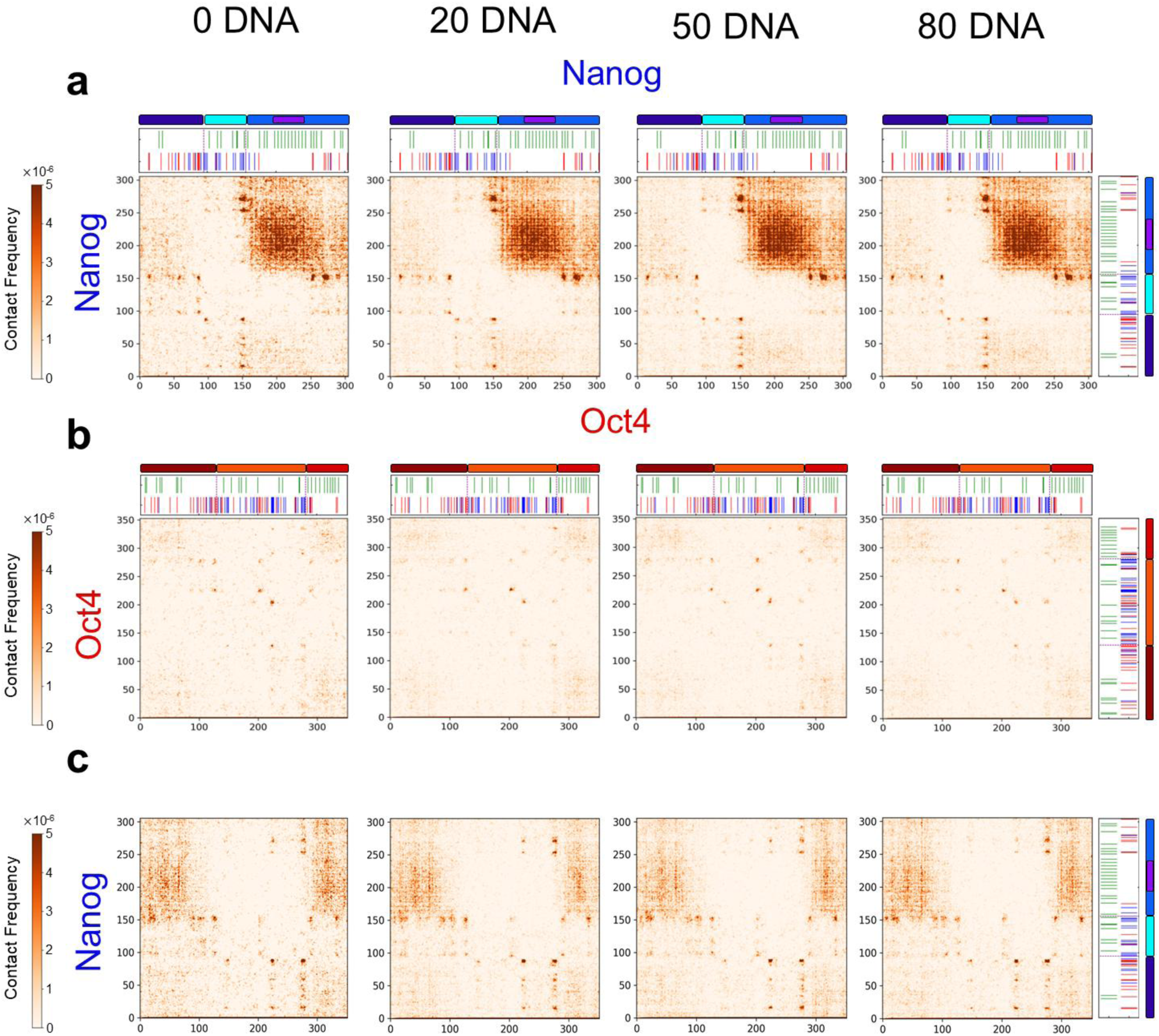
: DNA concentration–dependent protein–protein interactions in Nanog–Oct4 condensates. Residue-level contact maps computed from phase-separated Nanog-Oct4 mixtures at increasing DNA concentrations (0, 20, 50, and 80 DNA molecules; left to right). **(a)** Nanog–Nanog contact maps. Domain architecture and sequence chemical features of Nanog are shown above and to the right of each map. **(b)** Oct4-Oct4 contact maps under the same DNA conditions. Structural domains and sequence features of Oct4 are indicated above and to the right. **(c)** Nanog-Oct4 heterotypic contact maps across the four DNA concentrations. Domain and sequence annotations of Nanog are shown on the right following the same color code as in Fig. 1.

**Supplementary Figure 3.**
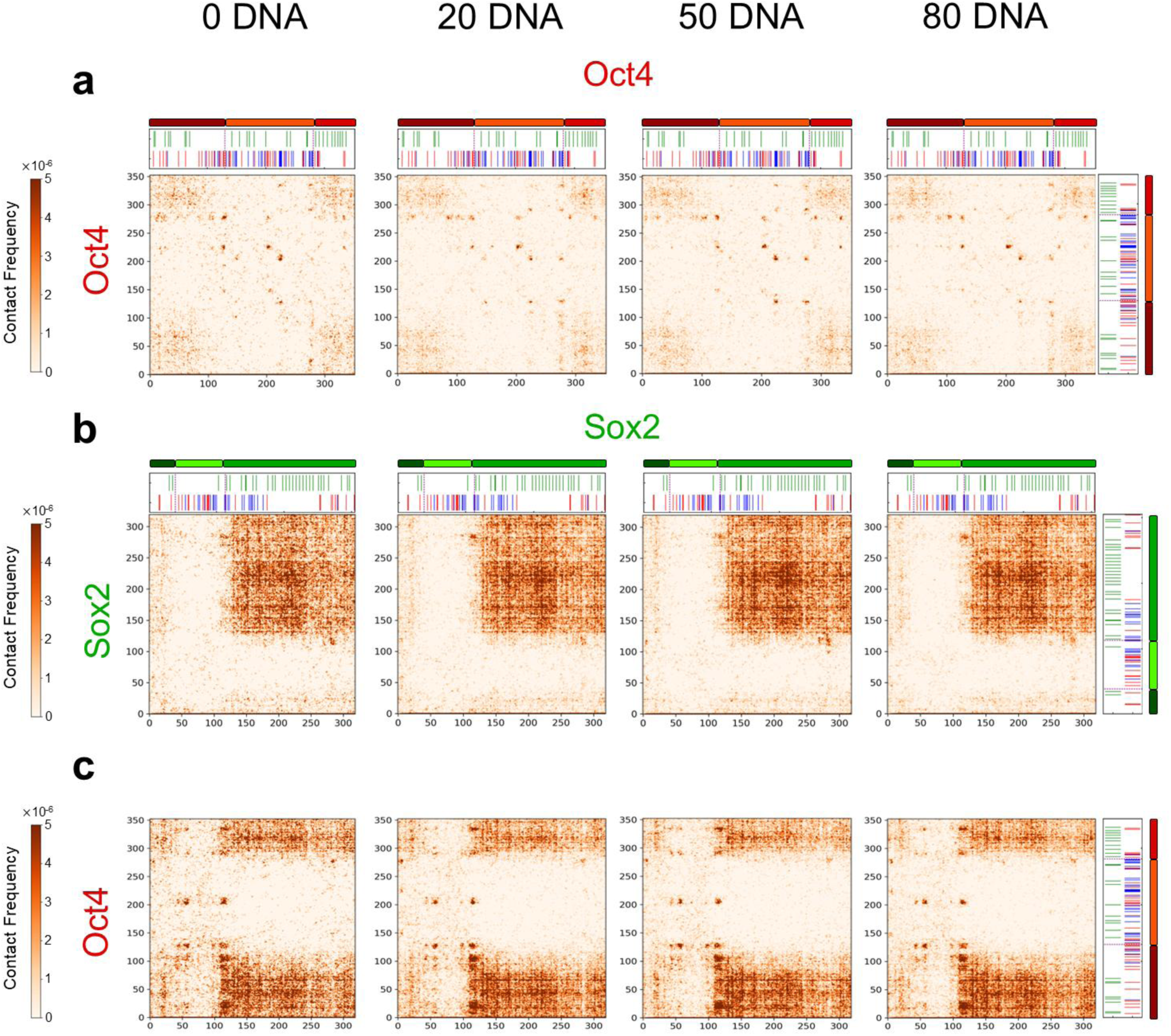
: DNA concentration–dependent interaction patterns in Oct4–Sox2 condensates. Residue-level contact maps computed from phase-separated Oct4-Sox2 mixtures at increasing DNA concentrations (0, 20, 50, and 80 DNA molecules; left to right). **(a)** Oct4-Oct4 homotypic contact maps. Domain architecture and sequence chemical features of Oct4 are shown above and to the right of each map. **(b)** Sox2-Sox2 homotypic contact maps under the same DNA conditions. Structural domains and sequence features of Sox2 are displayed above and to the right. **(c)** Oct4-Sox2 heterotypic contact maps across the four DNA concentrations. Domain and sequence annotations of Oct4 are shown on the right following the same color code as in Fig. 1.

**Supplementary Figure 4.**
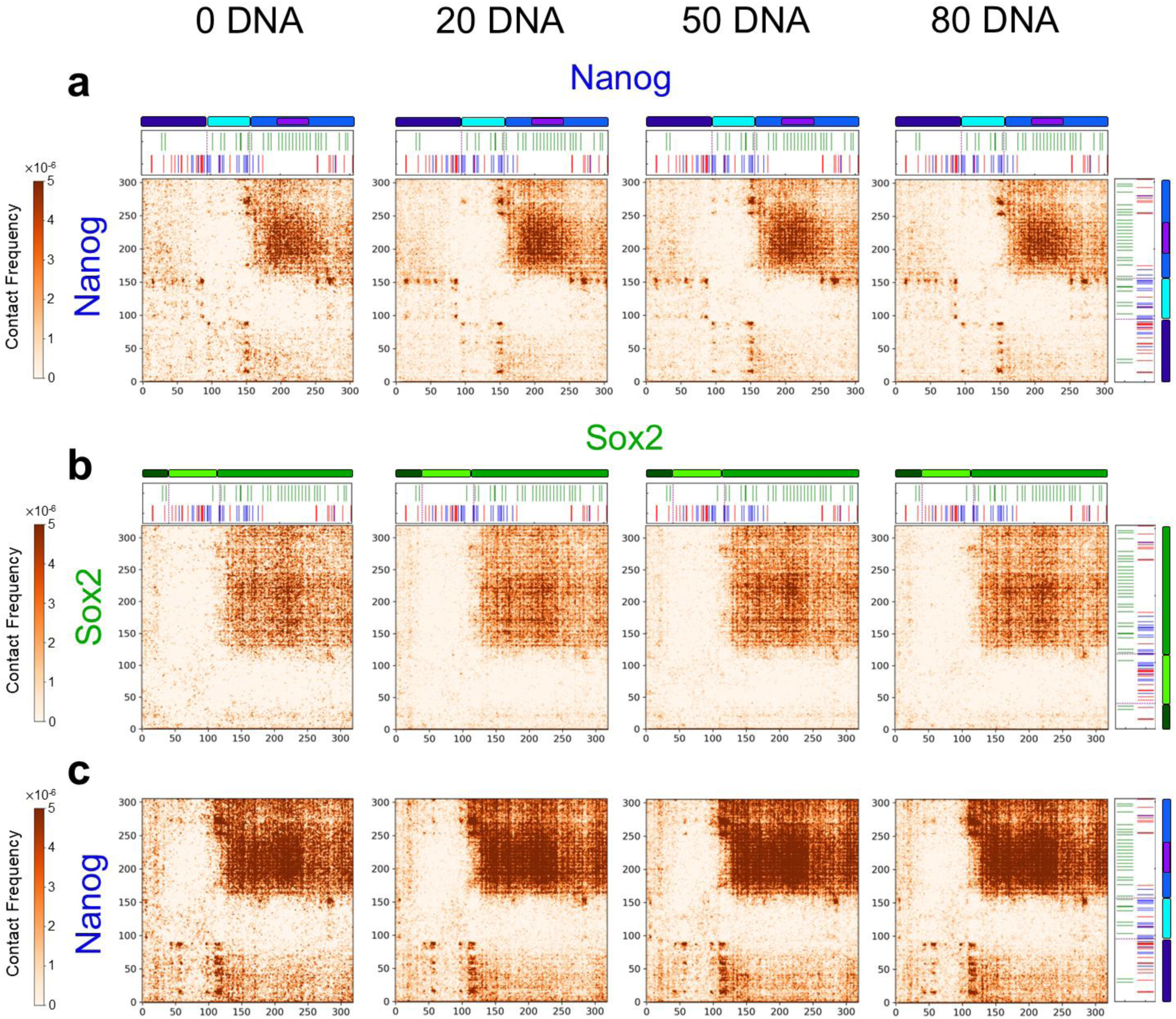
: DNA concentration-dependent interaction patterns in Nanog-Sox2 condensates. Residue-level contact maps computed from phase-separated Nanog-Sox2 mixtures at increasing DNA concentrations (0, 20, 50, and 80 DNA molecules; left to right). **(a)** Nanog-Nanog homotypic contact maps. Domain architecture and sequence chemical features of Nanog are shown above and to the right of each map. **(b)** Sox2-Sox2 homotypic contact maps under the same DNA conditions. Structural domains and sequence features of Sox2 are displayed above and to the right. **(c)** Nanog-Sox2 heterotypic contact maps across the four DNA concentrations. Domain and sequence annotations of Nanog are shown on the right following the same color code as in Fig. 1.

**Supplementary Figure 5.**
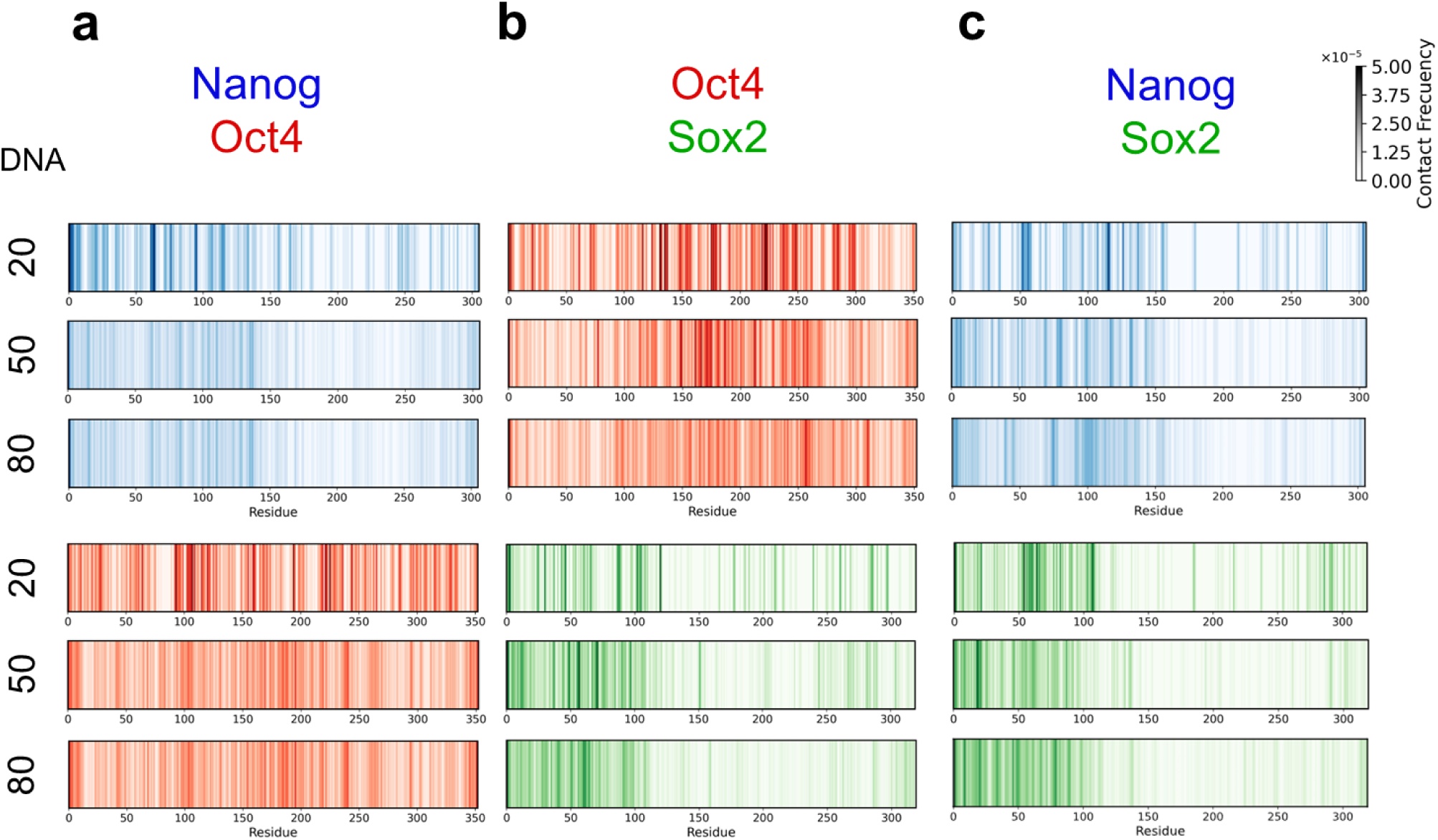
: DNA-protein contact patterns in phase-separated TF condensates at increasing DNA concentrations. Residue-resolved DNA-protein contact frequencies computed from binary condensates at 20, 50, and 80 DNA chains (indicated on the left). Color intensity reflects contact frequency (scale shown on the right). **(a)** Nanog-Oct4 mixtures: DNA-Nanog (blue) and DNA-Oct4 (red) contact maps. **(b)** Oct4-Sox2 mixtures: DNA-Oct4 (red) and DNA-Sox2 (green) contact maps. **(c)** Nanog-Sox2 mixtures: DNA-Nanog (blue) and DNA-Sox2 (green) contact maps.

**Supplementary Figure 6.**
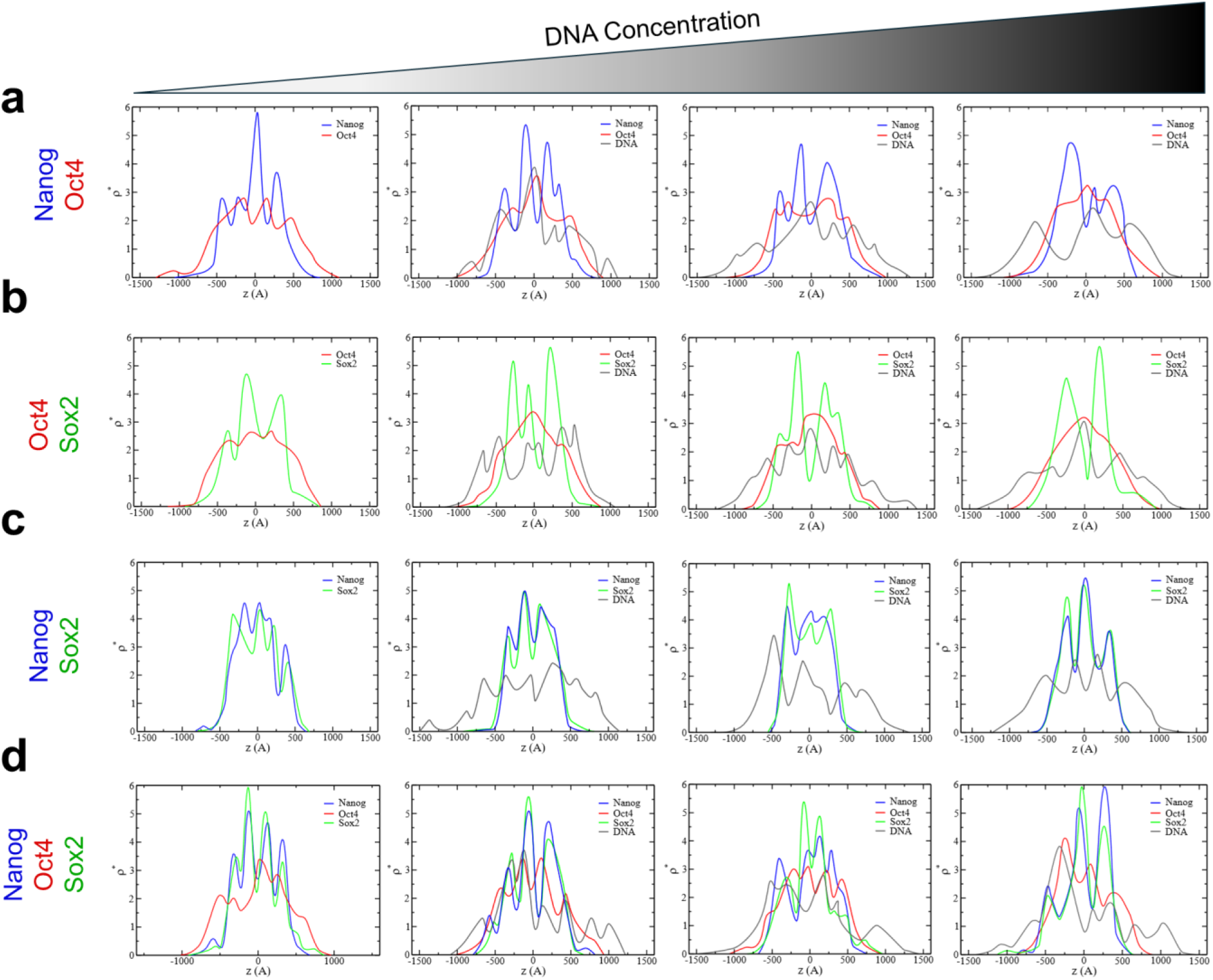
: Density profiles of phase-separated transcriptional condensates at increasing DNA concentrations. One-dimensional density profiles along the simulation box axis are shown for increasing DNA concentrations (0, 20, 50, and 80 DNA molecules; left to right). Protein species are color-coded as Nanog (blue), Oct4 (red), and Sox2 (green); DNA is shown in gray. **(a)** Nanog-Oct4 mixtures. **(b)** Oct4-Sox2 mixtures. **(c)** Nanog-Sox2 mixtures. **(d)** Nanog-Oct4-Sox2 ternary mixtures.

**Supplementary Figure 7.**
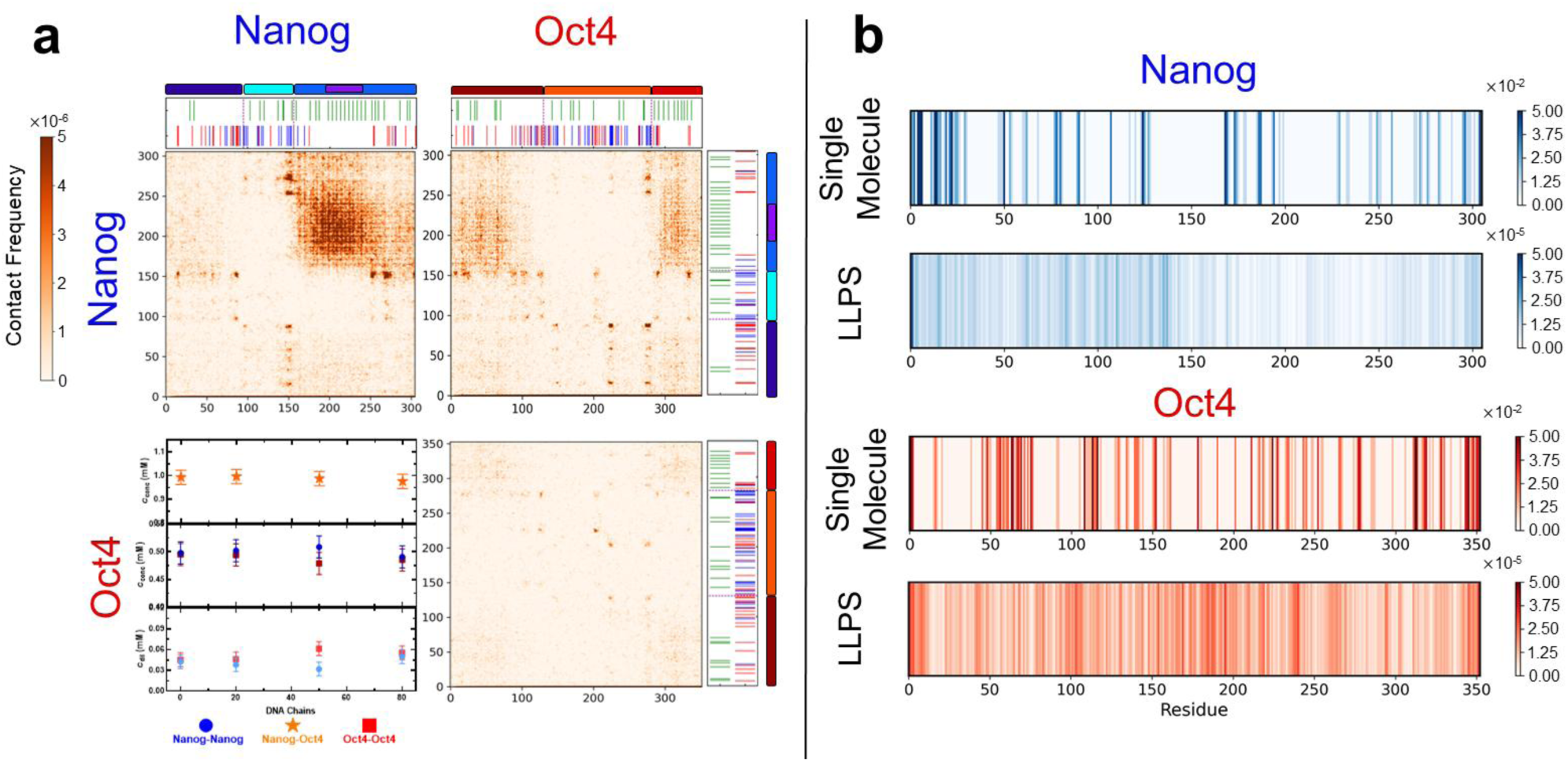
: Interaction patterns in DNA-containing Nanog-Oct4 condensates. **(a)** Residue-level protein-protein contact maps obtained from simulations of a Nanog-Oct4 mixture containing 100 Nanog, 100 Oct4, and 80 DNA chains. Structural domains and sequence chemical features are displayed above and to the right of the maps following the same color code as in Fig. 1. Top left: Nanog-Nanog; top right: Nanog-Oct4; bottom right: Oct4-Oct4 interactions. Bottom left: critical concentrations of each component as a function of DNA chain number, shown for the dilute phase (bottom), condensed phase (center), and total system (top). **(b)** Comparison of DNA-protein contact patterns between a single-molecule simulation (1 Nanog, 1 Oct4, 1 DNA) and a phase-separated system (100 Nanog, 100 Oct4, 80 DNA). DNA-Nanog interactions are shown in blue and DNA-Oct4 interactions in red.

**Supplementary Figure 8.**
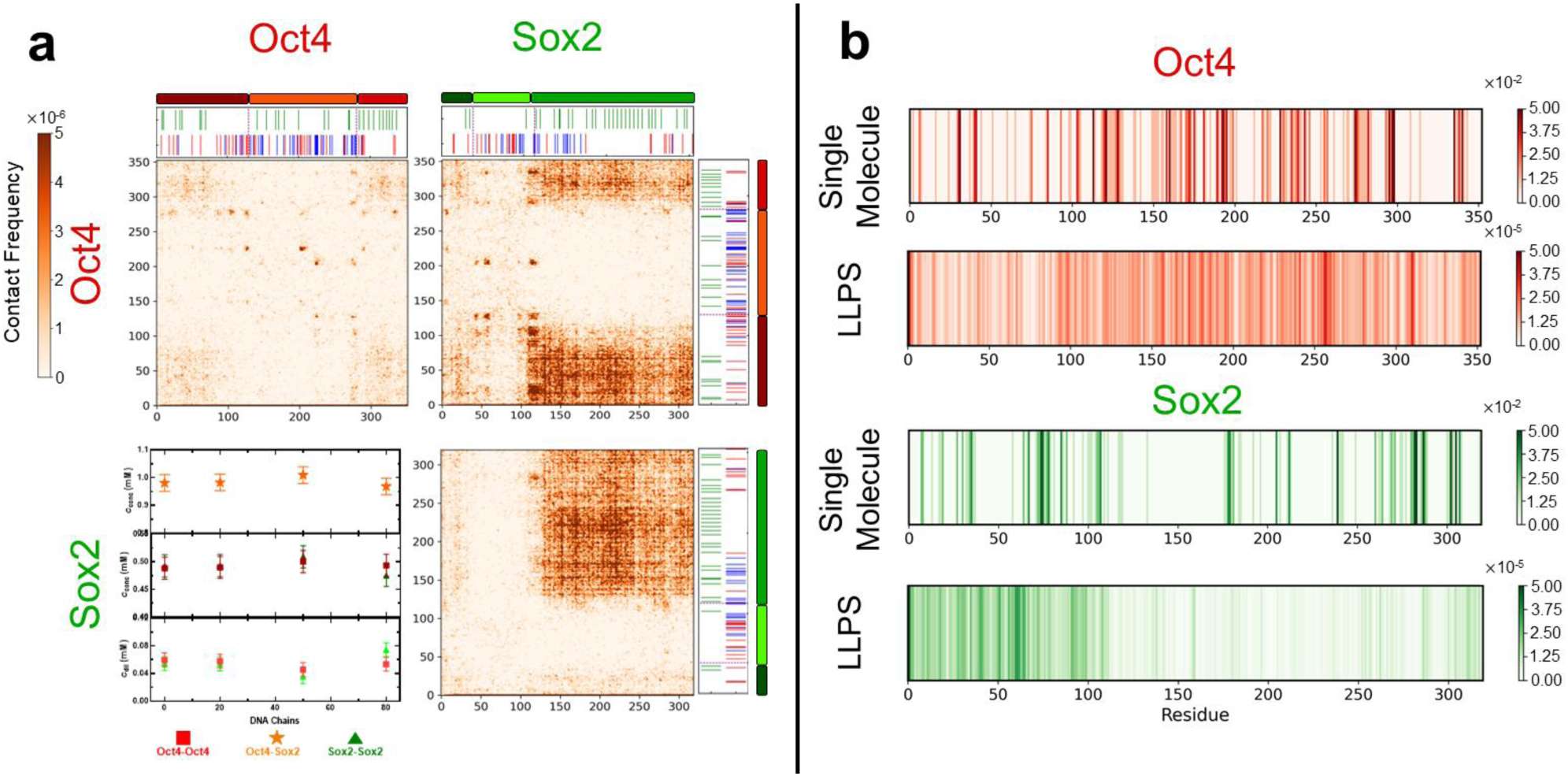
: Interaction patterns in DNA-containing Oct4-Sox2 condensates. **(a)** Residue-level protein-protein contact maps from simulations of a phase-separated mixture containing 100 Oct4, 100 Sox2, and 80 DNA chains. Structural domains and sequence chemical features are shown above and to the right of each map following the same color code as in Fig. 1. Top left: Oct4-Oct4; top right: Oct4-Sox2; bottom right: Sox2-Sox2 interactions. Bottom left: critical concentrations of each component as a function of DNA chain number, shown for the dilute phase (bottom), condensed phase (center), and the total system (top). **(b)** Comparison of DNA-protein contact patterns between a single-molecule simulation (1 Oct4, 1 Sox2, 1 DNA) and a condensed-phase simulation (100 Oct4, 100 Sox2, 80 DNA). DNA-Oct4 interactions are shown in red and DNA-Sox2 interactions in green.

**Supplementary Figure 9.**
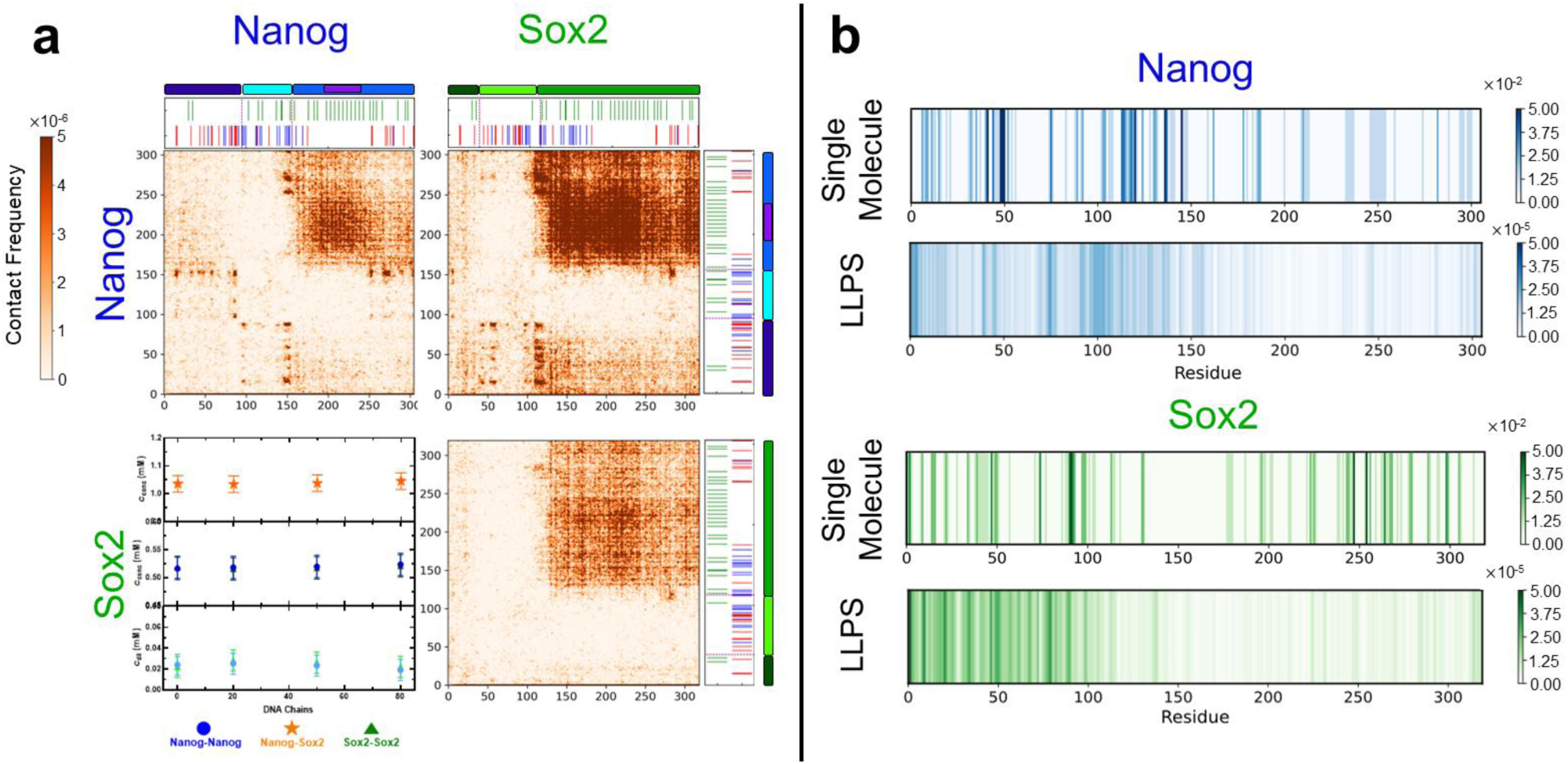
: Interaction patterns in DNA-containing Nanog-Sox2 condensates. **(a)** Residue-level protein-protein contact maps from simulations of a phase-separated mixture containing 100 Nanog, 100 Sox2, and 80 DNA chains. Structural domains and sequence chemical features are shown above and to the right of each map following the same color code as in Fig. 1. Top left: Nanog-Nanog; top right: Nanog-Sox2; bottom right: Sox2-Sox2 interactions. Bottom left: critical concentrations of each component as a function of DNA chain number, shown for the dilute phase (bottom), condensed phase (center), and the total system (top). **(b)** Comparison of DNA-protein contact patterns between a single-molecule simulation (1 Nanog, 1 Sox2, 1 DNA) and a condensed-phase simulation (100 Nanog, 100 Sox2, 80 DNA). DNA-Nanog interactions are shown in blue and DNA-Sox2 interactions in green.

## Description of Additional Supplementary Files

Supplementary Movie 1: Formation of a phase-separated transcriptional condensate formed by the ternary mixture of Nanog, Oct4 and Sox2. Representative simulation trajectory showing Nanog, Oct4, and Sox2 assembling into a phase-separated transcriptional condensate.

Supplementary Movie 2: Formation of a phase-separated transcriptional condensate formed by the binary mixture of Nanog and Oct4. Representative simulation trajectory showing Nanog and Oct4 assembling into a phase-separated transcriptional condensate.

Supplementary Movie 3: Formation of a phase-separated transcriptional condensate formed by the binary mixture of Oct4 and Sox2. Representative simulation trajectory showing Oct4 and Sox2 assembling into a phase-separated transcriptional condensate.

Supplementary Movie 4: Formation of a phase-separated transcriptional condensate formed by the binary mixture of Nanog and Sox2. Representative simulation trajectory showing Nanog and Sox2 assembling into a phase-separated transcriptional condensate.

**Supplementary Movie 5: DNA-mediated formation of a phase-separated transcriptional condensate.** Representative simulation trajectory of Nanog, Oct4, and Sox2 forming a phase-separated transcriptional condensate in the presence of 80 DNA chains.

## REFERENCE

1. Sabari, B. R. et al. Coactivator condensation at super-enhancers links phase separation and gene control. Science 361, eaar3958 (2018).

2. Boija, A. et al. Transcription Factors Activate Genes through the Phase-Separation Capacity of Their Activation Domains. Cell 175, 1842–1855.e16 (2018).

3. Guo, Y. E. et al. Pol II phosphorylation regulates a switch between transcriptional and splicing condensates. Nature 572, 543–548 (2019).

4. Zuo, L. et al. Loci-specific phase separation of FET fusion oncoproteins promotes gene transcription. Nat Commun 12, 1491 (2021).

5. Wei, M.-T. et al. Nucleated transcriptional condensates amplify gene expression. Nat Cell Biol 22, 1187–1196 (2020).

6. Hnisz, D., Shrinivas, K., Young, R. A., Chakraborty, A. K. & Sharp, P. A. A Phase Separation Model for Transcriptional Control. Cell 169, 13–23 (2017).

7. Zhou, Q., Chipperfield, H., Melton, D. A. & Wong, W. H. A gene regulatory network in mouse embryonic stem cells. Proceedings of the National Academy of Sciences 104, 16438–16443 (2007).

8. Takahashi, K. & Yamanaka, S. Induction of Pluripotent Stem Cells from Mouse Embryonic and Adult Fibroblast Cultures by Defined Factors. Cell 126, 663–676 (2006).

9. Rodda, D. J. et al. Transcriptional regulation of nanog by OCT4 and SOX2. J Biol Chem 280, 24731–24737 (2005).

10. Chambers, I. & Tomlinson, S. R. The transcriptional foundation of pluripotency. Development 136, 2311–2322 (2009).

11. Rizzino, A. Concise Review: The Sox2-Oct4 Connection: Critical Players in a Much Larger Interdependent Network Integrated at Multiple Levels. Stem Cells 31, 1033–1039 (2013).

12. Molliex, A. et al. Phase Separation by Low Complexity Domains Promotes Stress Granule Assembly and Drives Pathological Fibrillization. Cell 163, 123–133 (2015).

13. Wang, J. et al. A Molecular Grammar Governing the Driving Forces for Phase Separation of Prion-like RNA Binding Proteins. Cell 174, 688–699.e16 (2018).

14. Nguyen, T. et al. Chromatin sequesters pioneer transcription factor Sox2 from exerting force on DNA. Nat Commun 13, 3988 (2022).

15. Wang, J. et al. Phase separation of OCT4 controls TAD reorganization to promote cell fate transitions. Cell Stem Cell 28, 1868–1883.e11 (2021).

16. Huertas, J., Maristany, M. J. & Collepardo-Guevara, R. Oct4 clusters promote DNA accessibility by enhancing chromatin plasticity. Preprint at 10.1101/2025.10.20.683403.

17. Choi, K.-J. et al. NANOG prion-like assembly mediates DNA bridging to facilitate chromatin reorganization and activation of pluripotency. Nat Cell Biol 24, 737–747 (2022).

18. Mizutani, A., Tan, C., Sugita, Y. & Takada, S. Micelle-like clusters in phase-separated Nanog condensates: A molecular simulation study. PLOS Computational Biology 19, e1011321 (2023).

19. Whyte, W. A. et al. Master Transcription Factors and Mediator Establish Super-Enhancers at Key Cell Identity Genes. Cell 153, 307–319 (2013).

20. Farag, M., Borcherds, W. M., Bremer, A., Mittag, T. & Pappu, R. V. Phase separation of protein mixtures is driven by the interplay of homotypic and heterotypic interactions. Nat Commun 14, 5527 (2023).

21. Driver, M. D. & Onck, P. R. Selective phase separation of transcription factors is driven by orthogonal molecular grammar. Nat Commun 16, 3087 (2025).

22. Galvanetto, N. et al. Extreme dynamics in a biomolecular condensate. Nature 619, 876–883 (2023).

23. Dignon, G. L., Zheng, W., Kim, Y. C., Best, R. B. & Mittal, J. Sequence determinants of protein phase behavior from a coarse-grained model. PLOS Computational Biology 14, e1005941 (2018).

24. Joseph, J. A. et al. Physics-driven coarse-grained model for biomolecular phase separation with near-quantitative accuracy. Nat Comput Sci 1, 732–743 (2021).

25. Tesei, G., Schulze, T. K., Crehuet, R. & Lindorff-Larsen, K. Accurate model of liquid–liquid phase behavior of intrinsically disordered proteins from optimization of single-chain properties. Proceedings of the National Academy of Sciences 118, e2111696118 (2021).

26. Thomasen, F. E. et al. Rescaling protein-protein interactions improves Martini 3 for flexible proteins in solution. Nat Commun 15, 6645 (2024).

27. Li, C. et al. Deciphering the molecular mechanism underlying morphology transition in two-component DNA-protein cophase separation. Structure 33, 62–77.e8 (2025).

28. Bian, Y. et al. Structural organization of HBV pgRNA genome driven by phase separation in capsid confinement. Nat Commun 10.1038/s41467-026-69689-2 (2026) doi: 10.1038/s41467-026-69689-2.

29. Regy, R. M., Zheng, W. & Mittal, J. Using a sequence-specific coarse-grained model for studying protein liquid–liquid phase separation. in Methods in Enzymology vol. 646 1–17 (Academic Press, 2021).

30. R. Tejedor, A., et al. Chemically Informed Coarse-Graining of Electrostatic Forces in Charge-Rich Biomolecular Condensates. ACS Cent. Sci. 11, 302–321 (2025).

31. Li, W., Wang, W. & Takada, S. Energy landscape views for interplays among folding, binding, and allostery of calmodulin domains. Proceedings of the National Academy of Sciences 111, 10550–10555 (2014).

32. Mizutani, A., Tan, C., Sugita, Y. & Takada, S. Heterogeneous condensates of transcription factors in embryonic stem cells: Molecular simulations. Biophysical Journal 124, 1587–1598 (2025).

33. Knotts, T. A., IV, Rathore, N., Schwartz, D. C. & de Pablo, J. J. A coarse grain model for DNA. J. Chem. Phys. 126, 084901 (2007).

34. Freeman, G. S., Hinckley, D. M., Lequieu, J. P., Whitmer, J. K. & de Pablo, J. J. Coarse-grained modeling of DNA curvature. J. Chem. Phys. 141, 165103 (2014).

35. Terakawa, T., Kenzaki, H. & Takada, S. p53 Searches on DNA by Rotation-Uncoupled Sliding at C-Terminal Tails and Restricted Hopping of Core Domains. J. Am. Chem. Soc. 134, 14555–14562 (2012).

36. Tan, C., Terakawa, T. & Takada, S. Dynamic Coupling among Protein Binding, Sliding, and DNA Bending Revealed by Molecular Dynamics. J. Am. Chem. Soc. 138, 8512–8522 (2016).

37. Nagae, F., Brandani, G. B., Takada, S. & Terakawa, T. The lane-switch mechanism for nucleosome repositioning by DNA translocase. Nucleic Acids Res 49, 9066–9076 (2021).

38. Nagae, F., Takada, S. & Terakawa, T. Histone chaperone Nap1 dismantles an H2A/H2B dimer from a partially unwrapped nucleosome. Nucleic Acids Res 51, 5351–5363 (2023).

39. Nagae, F., Murayama, Y. & Terakawa, T. Molecular mechanism of parental H3/H4 recycling at a replication fork. Nat Commun 15, 9485 (2024).

40. Gu, C., Takada, S., Brandani, G. B. & Terakawa, T. Molecular dynamics simulations of human cohesin subunits identify DNA binding sites and their potential roles in DNA loop extrusion. PLOS Computational Biology 21, e1012493 (2025).

41. Nagae, F., Murata, Y., Yamauchi, M., Takada, S. & Terakawa, T. Mechanistic models of asymmetric hand-over-hand translocation and nucleosome navigation by CMG helicase. Nat Commun 16, 10304 (2025).

42. Murata, Y., Niina, T. & Takada, S. OpenCafeMol: A coarse-grained biomolecular simulator on GPU with its application to vesicle fusion. Biophysical Journal 125, 420–431 (2025).

43. Heurtier, V. et al. The molecular logic of Nanog-induced self-renewal in mouse embryonic stem cells. Nat Commun 10, 1109 (2019).

44. King, H. W. & Klose, R. J. The pioneer factor OCT4 requires the chromatin remodeller BRG1 to support gene regulatory element function in mouse embryonic stem cells. eLife 6, e22631 (2017).

45. Esch, D. et al. A unique Oct4 interface is crucial for reprogramming to pluripotency. Nat Cell Biol 15, 295–301 (2013).

46. Chen, X. et al. Integration of External Signaling Pathways with the Core Transcriptional Network in Embryonic Stem Cells. Cell 133, 1106–1117 (2008).

47. Hou, Y. et al. Emerging cooperativity between Oct4 and Sox2 governs the pluripotency network in early mouse embryos. eLife 13, RP100735 (2025).

48. Avilion, A. A. et al. Multipotent cell lineages in early mouse development depend on SOX2 function. Genes Dev 17, 126–140 (2003).

49. Novak, D. et al. SOX2 in development and cancer biology. Seminars in Cancer Biology 67, 74–82 (2020).

50. Zhang, S. & Cui, W. Sox2, a key factor in the regulation of pluripotency and neural differentiation. World J Stem Cells 6, 305–311 (2014).

51. Patel, A. et al. A Liquid-to-Solid Phase Transition of the ALS Protein FUS Accelerated by Disease Mutation. Cell 162, 1066–1077 (2015).

52. Ferrolino, M. C., Mitrea, D. M., Michael, J. R. & Kriwacki, R. W. Compositional adaptability in NPM1-SURF6 scaffolding networks enabled by dynamic switching of phase separation mechanisms. Nat Commun 9, 5064 (2018).

53. Jang, S. et al. Phosphofructokinase relocalizes into subcellular compartments with liquid-like properties in vivo. Biophys J 120, 1170–1186 (2021).

54. Erkamp, N. A. et al. Differential interactions determine anisotropies at interfaces of RNA-based biomolecular condensates. Nat Commun 16, 3463 (2025).

55. Garaizar, A. et al. Aging can transform single-component protein condensates into multiphase architectures. Proceedings of the National Academy of Sciences 119, e2119800119 (2022).

56. Farag, M. et al. Condensates formed by prion-like low-complexity domains have small-world network structures and interfaces defined by expanded conformations. Nat Commun 13, 7722 (2022).

57. Tejedor, A. R., Collepardo-Guevara, R., Ramírez, J. & Espinosa, J. R. Time-Dependent Material Properties of Aging Biomolecular Condensates from Different Viscoelasticity Measurements in Molecular Dynamics Simulations. J. Phys. Chem. B 127, 4441–4459 (2023).

58. Tejedor, A. R. et al. Protein structural transitions critically transform the network connectivity and viscoelasticity of RNA-binding protein condensates but RNA can prevent it. Nat Commun 13, 5717 (2022).

59. Krainer, G. et al. Reentrant liquid condensate phase of proteins is stabilized by hydrophobic and non-ionic interactions. Nat Commun 12, 1085 (2021).

60. Terakawa, T. & Takada, S. Multiscale Ensemble Modeling of Intrinsically Disordered Proteins: p53 N-Terminal Domain. Biophysical Journal 101, 1450–1458 (2011).

61. Tesei, G., Schulze, T. K., Crehuet, R. & Lindorff-Larsen, K. Accurate model of liquid–liquid phase behavior of intrinsically disordered proteins from optimization of single-chain properties. Proceedings of the National Academy of Sciences 118, e2111696118 (2021).

62. Das, S., Lin, Y.-H., Vernon, R. M., Forman-Kay, J. D. & Chan, H. S. Comparative roles of charge, π, and hydrophobic interactions in sequence-dependent phase separation of intrinsically disordered proteins. Proceedings of the National Academy of Sciences 117, 28795–28805 (2020).

63. Wang, J. et al. A Molecular Grammar Governing the Driving Forces for Phase Separation of Prion-like RNA Binding Proteins. Cell 174, 688–699.e16 (2018).

64. Rana, U. et al. Asymmetric oligomerization state and sequence patterning can tune multiphase condensate miscibility. Nat. Chem. 16, 1073–1082 (2024).

65. Huangfu, D. et al. Induction of pluripotent stem cells from primary human fibroblasts with only Oct4 and Sox2. Nat Biotechnol 26, 1269–1275 (2008).

66. Malaga Gadea, F. C. & Nikolova, E. N. Structural Plasticity of Pioneer Factor Sox2 and DNA Bendability Modulate Nucleosome Engagement and Sox2-Oct4 Synergism. Journal of Molecular Biology 435, 167916 (2023).

67. Wagh, K., Garcia, D. A. & Upadhyaya, A. Phase separation in transcription factor dynamics and chromatin organization. Current Opinion in Structural Biology 71, 148–155 (2021).

68. Michael, A. K. et al. Mechanisms of OCT4-SOX2 motif readout on nucleosomes. Science 368, 1460–1465 (2020).

69. Borcherds, W., Bremer, A., Borgia, M. B. & Mittag, T. How do intrinsically disordered protein regions encode a driving force for liquid–liquid phase separation? Current Opinion in Structural Biology 67, 41–50 (2021).

70. Banani, S. F., Lee, H. O., Hyman, A. A. & Rosen, M. K. Biomolecular condensates: organizers of cellular biochemistry. Nat Rev Mol Cell Biol 18, 285–298 (2017).

71. Echigoya, K. et al. Nucleosome binding by the pioneer transcription factor OCT4. Sci Rep 10, 11832 (2020).

72. Nocente, M. C. et al. cBAF generates subnucleosomes that expand OCT4 binding and function beyond DNA motifs at enhancers. Nat Struct Mol Biol 31, 1756–1768 (2024).

73. Alston, H., Rouches, M., Murugan, A., Walczak, A. M. & Mora, T. Theoretical limits for sensing through phase separation. Proceedings of the National Academy of Sciences 123, e2520040123 (2026).

74. Schuster, B. S. et al. Identifying sequence perturbations to an intrinsically disordered protein that determine its phase-separation behavior. Proceedings of the National Academy of Sciences 117, 11421–11431 (2020).

75. Dannenhoffer-Lafage, T. & Best, R. B. A Data-Driven Hydrophobicity Scale for Predicting Liquid–Liquid Phase Separation of Proteins. J. Phys. Chem. B 125, 4046–4056 (2021).

76. Feito, A. et al. Benchmarking residue-resolution protein coarse-grained models for simulations of biomolecular condensates. PLOS Computational Biology 21, e1012737 (2025).

77. Blazquez, S. et al. Location and Concentration of Aromatic-Rich Segments Dictates the Percolating Inter-Molecular Network and Viscoelastic Properties of Ageing Condensates. Advanced Science 10, 2207742 (2023).

78. Niina, T., Brandani, G. B., Tan, C. & Takada, S. Sequence-dependent nucleosome sliding in rotation-coupled and uncoupled modes revealed by molecular simulations. PLOS Computational Biology 13, e1005880 (2017).

79. Brandani, G. B., Tan, C. & Takada, S. The kinetic landscape of nucleosome assembly: A coarse-grained molecular dynamics study. PLOS Computational Biology 17, e1009253 (2021).

80. Fiser, A. & Sali, A. Modeller: generation and refinement of homology-based protein structure models. Methods Enzymol 374, 461–491 (2003).

81. Jauch, R., Ng, C. K. L., Saikatendu, K. S., Stevens, R. C. & Kolatkar, P. R. Crystal Structure and DNA Binding of the Homeodomain of the Stem Cell Transcription Factor Nanog. Journal of Molecular Biology 376, 758–770 (2008).

82. Jumper, J. et al. Highly accurate protein structure prediction with AlphaFold. Nature 596, 583–589 (2021).

83. Lu, X.-J. & Olson, W. K. 3DNA: a versatile, integrated software system for the analysis, rebuilding and visualization of three-dimensional nucleic-acid structures. Nat Protoc 3, 1213–1227 (2008).

84. Michaud-Agrawal, N., Denning, E. J., Woolf, T. B. & Beckstein, O. MDAnalysis: A toolkit for the analysis of molecular dynamics simulations. Journal of Computational Chemistry 32, 2319–2327 (2011).

